# TLR2 signaling uniquely destabilizes tumor Tregs to promote cancer immunotherapy

**DOI:** 10.1101/2025.06.26.661786

**Authors:** Timothy F. Campbell, Jesse Garcia Castillo, Sebastian Fernandez, Diego Gonzalez Ventura, Jenna Vickery, Stefan Homentcovschi, Nicole Flores Hernandez, Julia Ybarra, Daniel A. Portnoy, Michel DuPage

**Affiliations:** Division of Immunology and Molecular Medicine, Department of Molecular and Cell Biology, University of California, Berkeley, Berkeley, CA 94720, USA; Department of Plant and Microbiology, University of California, Berkeley, Berkeley, CA 94720, USA

**Author notes:** Contact info.

## Abstract

A fundamental principle of immune responses is that innate immunity promotes adaptive immunity, but in the context of cancer immunity, the importance of innate receptor signaling is poorly defined. To study how Toll-like receptor (TLR) signaling contributes to cancer immunity, we used an attenuated strain of Listeria monocytogenes (Lm) that inhibits tumor growth in mice. We found that Lm stimulation of TLR2, but not TLR5 or TLR9, was essential for tumor control. As expected, TLR2 supported the priming of Lm-specific T cells in lymphoid tissues. However, TLR2 signaling in innate immune cells within tumors also destabilized Foxp3 expression in regulatory T cells (Tregs), a function that was not mediated by other TLRs. Reduced Treg function promoted tumor antigen cross-presentation by DC1s to enhance the functionality of recalled tumor-specific CD8+ T cells directly within tumors. These findings reveal a unique capacity for TLR2 signaling to diminish immunosuppression and promote cancer immunity.

## Introduction

Recent advancements in cancer immunotherapy have focused on targeting T cells that kill cancer cells.^1,2^ Therapeutic cancer vaccination, which uses microbial vectors like attenuated *Listeria monocytogenes* (*Lm*) engineered to express tumor antigens, is one such strategy that aims to elicit antitumor T cell mediated immunity.^3,4^ While bacterial therapeutic vaccines have shown promise in preclinical models, most have not demonstrated sufficient efficacy in clinical trials.^3,4^ An advantage of microbial cancer vaccines is that pattern recognition receptor (PRR) ligands produced by microbial vectors promote tumor-specific T cell priming, allowing a single agent to serve as both antigen and adjuvant.^3,4^ However, beyond their role in T cell priming, the effects of PRR stimulation by cancer vaccine vectors remain poorly understood.^5,6^ Furthermore, in vectors like *Lm* that activate multiple PRRs, it is unclear whether a single PRR ligand provides unique adjuvant function or whether redundant pathways collectively drive antitumor efficacy.^7^ The failure of prior clinical trials suggests that successful therapeutic cancer vaccination will require not only optimizing tumor-specific T cell priming but also taking full advantage of PRR signaling to enhance antitumor immunity.

Toll-like receptors (TLRs), a well-characterized PRR family, are among the first innate receptors to be engaged during infection.^8^ TLR signaling in antigen-presenting cells (APCs) such as dendritic cells (DCs) upregulates genes involved in antigen presentation and costimulation that are crucial for initiating T cell-mediated immune responses.^8^ TLRs and APCs thus play key roles in shaping antitumor T cell responses elicited by cancer vaccines.^8–12^ Accordingly, intratumoral injection of TLR ligands can inhibit tumor growth in mice, presumably by increasing T cell priming against tumor antigens.^9,10,13^ Clinical testing of TLR ligands has emphasized the viral-sensing endosomal TLRs (TLR3, 7/8, and 9), leading to the approval of the TLR7 agonist imiquimod for treating superficial basal cell carcinoma.^10,13–17^ However, how TLR signaling affects antitumor immunity beyond its role in *de novo* T cell priming is poorly understood.^5,6^ Moreover, whether bacteria-sensing cell-surface TLRs (TLR1/2, 2/6, 4, and 5) can promote antitumor CD8^+^ T cell responses similarly to endosomal TLRs remains underexplored.

Several findings provide evidence that bacterially-mediated cell-surface TLR stimulation within tumors can benefit cancer immunotherapy. Intravesical BCG, which activates TLR2 and 4, is a standard treatment for noninvasive bladder cancer.^18^ Similarly, the first recorded form of cancer immunotherapy, Coley’s Toxins, employed direct injection of heat-killed *Streptococcus pyogenes* and *Serratia marcescens* (activating TLR2, TLR4, and TLR5) into sarcomas.^19–23^ *Lm* can likewise stimulate TLR2, 5, and 9, and TLR signaling is essential for controlling *L. monocytogenes* in non-tumor tissues, prompting speculation that TLR signaling by *Lm* cancer vaccines is beneficial.^9,24^ However, studies of anti-*Lm* immunity outside the context of cancer vaccination suggest that anti-*Lm* CD8^+^ T cell immunity during secondary infection is largely TLR-independent.^7,25–31^ Likewise, TLR signaling-deficient patients, while more susceptible to bacterial infections early in life, appear to develop effective adaptive immunity to bacteria late in life, implying a limited role for TLRs beyond the control of primary infections and *de novo* T cell priming.^32,33^ Consequently, the effects of *Lm*-mediated TLR signaling on antitumor T cell responses are understudied, leaving several fundamental questions about the role of TLR signaling by *Lm* cancer vaccines unanswered. One open question is whether TLR signaling by *Lm* cancer vaccines modulates tumor antigen presentation. While *Lm* cancer vaccines deliver tumor antigens to the MHC-I presentation pathway upon entry into the cytosol, their effect on the presentation of unlinked, endogenous tumor antigens (i.e., cross-presentation) remains unclear.^3,4,7^ In steady-state conditions, tumor antigen cross-presentation by Batf3^+^ cDC1s is the primary mechanism of priming antitumor CD8^+^ T cells.^34,35^ Intriguingly, cross-presentation is also required to prime *Lm*-specific CD8^+^ T cells despite the ready access of *Lm* antigens to the endogenous MHC-I pathway.^36–38^ We recently found that anti-*Lm* T cell responses in tumors can facilitate the cross-presentation of tumor antigens.^39^ *Lm*-mediated TLR signaling within tumors may underlie this effect.^40,41^ Indeed, TLR ligands linked to tumor antigens can enhance cross-presentation and bolster antitumor T cell responses *in vivo*.^41,42^ However, whether TLR signaling regulates *Lm*-mediated cross-presentation - and if this affects endogenous tumor antigens in addition to *Lm* antigens - remains unknown.

Another unanswered question is how TLR stimulation by *Lm* affects immune suppression within the tumor microenvironment (TME), in particular the frequency and functionality of regulatory T cells (Tregs), which are a major impediment to CD8^+^ T cell-mediated antitumor immunity.^43^ Tregs have been shown to inhibit antitumor T cell responses by limiting tumor antigen cross-presentation,^44,45^ preferentially binding the T cell mitogenic cytokine IL-2,^46^ and expression of immunosuppressive genes such as CTLA-4, IL-10, and TGF-β.^47^ Tregs have been reported to be inhibited both directly and indirectly by TLR signaling,^48–50^ and infection of mice with *L. monocytogenes* and other microbes can reduce Treg frequency and inhibitory function.^51–54^ However, direct evidence to link reduced Treg function and frequency to TLR signaling by *L. monocytogenes* is lacking, and it is not known whether Treg modulation by TLRs plays a role in antitumor immunity in the context of *Lm*-based cancer immunotherapies.

Dual intravenous and then intratumoral *Lm* administration (IV+IT *Lm*) controls tumor growth in a Batf3^+^ cDC1- and CD8^+^ T cell-dependent manner by enhancing the CD8^+^ T cell response against *Lm* and tumor antigens.^39^ Here we show that TLR2 - but not TLR5 or TLR9 - is required for the efficacy of IV+IT *Lm* via three mechanisms. As expected, TLR2 supported the priming of *Lm*-specific CD8^+^ T cells during IV *Lm* administration. However, TLR2 signaling also exerted unanticipated effects during IT *Lm* administration that were distinct from its role in priming anti-*Lm* T cells. First, TLR2 contributed to the recall of *Lm*-specific CD8^+^ T cells to tumors following IT *Lm* administration. Second, TLR2 mediated tumor antigen cross-presentation by type I dendritic cells (DC1s), rapidly reversing tumor-specific CD8^+^ T cell exhaustion in tumors and improving *de novo* antitumor T cell priming in the tumor-draining lymph node. Third, TLR2 mediated the destabilization of Foxp3 expression in tumor-infiltrating Tregs, which reduced immune suppression within tumors and potentiated the antitumor effects of IV+IT *Lm*. Importantly, only TLR2 signaling, and not any other TLR tested, impacted Treg function in this manner, thus explaining the unique contribution of TLR2 during IT *Lm* administration. Together, these results demonstrate an unexpected role for TLR2 signaling in CD8^+^ T cell-mediated immunity during both primary and secondary immune responses that can be leveraged to enhance the efficacy of microbial cancer immunotherapies.

## Results

### IV+IT *Lm* immunotherapy requires TLR2 signaling for tumor control

To test the impact of TLR signaling on tumor control with IV+IT *Lm*, we performed the IV+IT therapeutic regimen in MC38 tumor-bearing mice that were deficient for each of the TLRs known to be stimulated by *Lm*, TLRs 2, 5, and 9, compared to WT controls (Figure 1A).^39,55^ We found that while *Tlr5* and *Tlr9* deficient mice were still able to control tumors, *Tlr2* deficient mice could not control tumors with IV+IT *Lm* (Figure 1B). Interestingly, tumor control in *Tlr2* deficient mice was already reduced by the day of IT *Lm* treatment (day 11, 7 days after IV *Lm*), suggesting that TLR2 signaling was important at the IV phase of *Lm* delivery (Figure S1A). Since anti-*Lm* CD8^+^ T cells were the primary mediators of tumor control in response to IV+IT *Lm* and are primed by IV *Lm* infection,^39^ we hypothesized that the loss of tumor control in *Tlr2*^-/-^ mice was due to a reduced CD8^+^ T cell response. Indeed, while CD8^+^ T cell frequency was increased in IV+IT *Lm*-treated WT, *Tlr5*^-/-^, and *Tlr9*^-/-^ mice, this effect was lost in *Tlr2*^-/-^ mice (Figure S1B). Accompanying the loss of tumor control was an increase in the CFU of *Lm* recovered from tumors in *Lm*-treated *Tlr2*^-/-^ mice compared to WT, *Tlr5*^-/-^, and *Tlr9*^-/-^ mice (Figure S1C).

**Figure 1:**
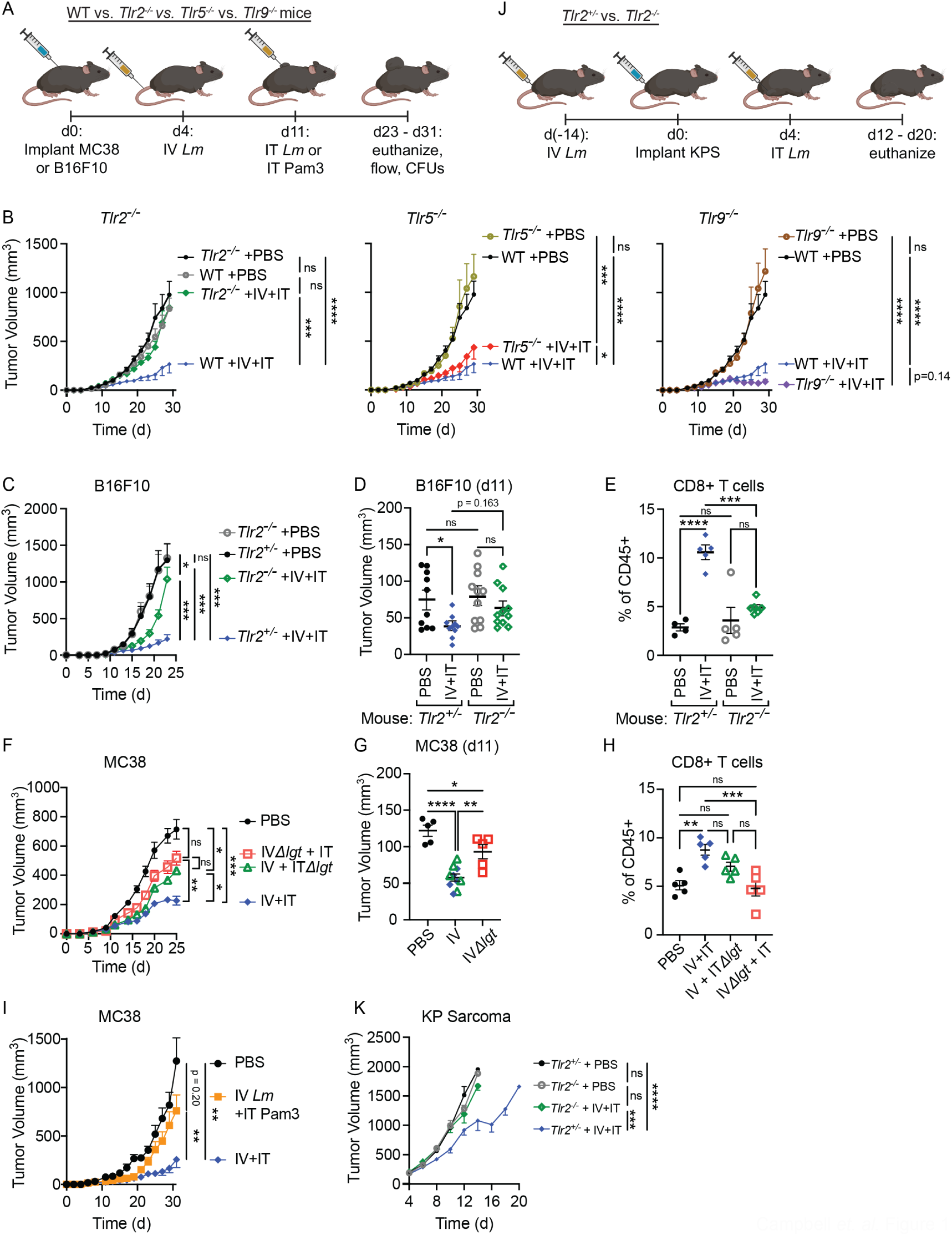
TLR2 signaling is required for the efficacy of IV+IT *Lm* therapy. (A) WT (*Tlr2^+/-^*), *Tlr2^-/^*^-^, *Tlr5^-/-^*, and *Tlr9^-/-^* mice were implanted subcutaneously with 5×10^5^ MC38 or B16F10 tumor cells (day 0). On day 4, mice received 1×10^6^ CFU IV *Lm*. On day 11, mice received 5×10^7^ CFU IT *Lm*. (B) MC38 tumor growth in WT (*Tlr2^+/-^*; PBS: n=10 mice; IV+IT *Lm*: n=8 mice), *Tlr2^- / -^* (n=9 mice/group)^-^, *Tlr5^-/-^* (PBS: n=8 mice; IV+IT *Lm*: n=10 mice), and *Tlr9^-/-^* (n=6 mice/group) mice receiving IV+IT *Lm* or PBS. (C) B16F10 tumor growth in *Tlr2^+/-^* (n=10 mice/group) and *Tlr2^-/-^* (n=11 mice/group) littermates receiving IV+IT *Lm* or PBS. (D) B16F10 tumor volume on day 11 of tumor growth from experiment in panel C. (E) CD8^+^ T cell frequencies in B16F10 tumors from experiment in panel C on day 23 of tumor growth (WT +PBS: n=4 mice; WT + IV+IT *Lm*: n=5 mice; *Tlr2^-/-^* + PBS: n=5 mice; *Tlr2^-/-^* + IV+IT *Lm*: n=6 mice). (F) MC38 tumor growth in WT mice receiving *LmΔlgt* during the IV stage (red squares), the IT stage (green triangles), or *lgt*-competent *Lm* at both stages of administration (blue diamonds) (n=5 mice/group). (G) MC38 tumor volumes on day 11 of tumor growth from experiment in panel F. (H) CD8^+^ T cell frequencies in MC38 tumors from experiment in panel F on day 25 of tumor growth. (I) Tumor growth in WT mice receiving IV *Lm* and IT PBS (n=10 mice) IT *Lm* (n=11 mice) or IT Pam3 (100 ug; n=9 mice). (J) *Tlr2^+/-^* and *Tlr2^-/-^* littermate mice received orthotopically implanted KP sarcomas. Two weeks prior to tumor growth, mice were treated with IV *Lm* (1×10^6^ CFU) or PBS (mock infection). 4 days after implantation, mice were treated with IT *Lm* (5×10^7^ CFU; n=12 mice/group) or PBS (n=12 mice/group). (K) KP sarcoma growth in *Tlr2^+/-^* and *Tlr2^-/-^* mice receiving IV+IT *Lm* or PBS. Mean ± s.e.m. and one-way ANOVA with multiple comparisons (D, E, G, H) or two-way ANOVA (B, C, F, I, K); P: *≤0.05, **≤0.01, ***≤0.001, ****≤ 0.000.

To determine whether the dependence of IV+IT *Lm* upon TLR2 signaling was applicable beyond the MC38 tumor model, we next treated *Tlr2*^+/-^ and *Tlr2*^-/-^ littermate mice bearing B16F10 tumors with IV+IT *Lm*. We again found that tumor control was significantly attenuated in *Tlr2*^-/-^ mice and that CD8^+^ T cell frequencies were reduced while *Lm* CFUs were increased in tumors of Tlr2-/- mice (Figures 1C-1E and S1D). More importantly, TLR2 was completely essential for the control of orthotopically implanted KrasG12D;p53-/- (KP) sarcomas with IV+IT *Lm* (Figure 1J-K and S1E).^56^ Together, these findings suggest that TLR2 plays a critical and conserved role in limiting *Lm* burden in tumor tissue and in mediating tumor control elicited by IV+IT *Lm* across multiple tumor types and anatomical locations.

### TLR2 signaling during IV and IT administration of *Lm* is required for tumor control

To test the requirement for TLR2 signaling specifically at the IV or IT stage of the IV+IT *Lm* regimen, we used an *LmΔlgt* strain, which does not generate the bacterial lipoprotein ligands for TLR2, during either IV or IT administration of *Lm* to MC38 tumor-bearing mice.^57,58^ Surprisingly, mice that were treated with *LmΔlgt* at either the IV or IT stage failed to fully control tumor growth with IV+IT *Lm* administration (Figure 1F). While IV *LmΔlgt* failed to slow tumor growth to the extent of IV *Lm* in the days preceding IT *Lm* administration (day 11, 7 days after IV *LmΔlgt*) (Figure 1G), substituting IT *LmΔlgt* after IV *Lm* also significantly reduced tumor control (Figure 1F). Thus, TLR2 signaling is necessary at both the IV and IT phases of *Lm* administration for full tumor control. Finally, as with *Tlr2*^-/-^ mice, *LmΔlgt* delivery at either the IV or IT phase led to reduced frequencies of tumor-infiltrating CD8^+^ T cells and more *Lm* CFUs in tumors (Figures 1H and S1F), suggesting that stimulation of TLR2 by *Lm* is important at both stages of the IV+IT *Lm* regimen.

### TLR2 signaling alone is not sufficient for tumor control with IV+IT *Lm*

To test whether TLR2 stimulation alone was sufficient to phenocopy IT *Lm* in the IV+IT *Lm* regimen, MC38 tumor-bearing mice were treated with IV *Lm* followed by either IT *Lm* or Pam3CSK4 (Pam3), a synthetic TLR2 ligand. IT Pam3 treatment initially controlled tumors but ultimately did not provide long-term control or clear *Lm* CFUs as compared to IT *Lm* (Figures 1I and S1G-H). Altogether, these results demonstrate that IT TLR2 signaling is necessary, but not sufficient, for tumor control and the clearance of *Lm* from tumors.

### TLR2 stimulation plays multiple roles in supporting CD8^+^ T cells in IV+IT *Lm* immunotherapy

Next, we sought to determine how TLR2 signaling impacted the CD8^+^T cell response at each stage of the IV+IT *Lm* regimen. We hypothesized that TLR2 signaling in antigen presenting cells could contribute to the expansion of *Lm*-specific CD8^+^ T cells during IV *Lm* administration. Therefore, we infected *Tlr2*^+/-^ and *Tlr2*^-/-^ littermate control mice with *Lm-OVA* and analyzed the frequency and activation of *Lm-OVA*-specific CD8^+^ T cells in the spleen using H-2K(b)-SIINFEKL tetramer staining (Figure 2A). The frequency of *Lm*-*OVA*- specific (tetramer+) CD8^+^ T cells and the activation of bulk CD8^+^ T cells (tetramer-) were reduced in *Tlr2*^-/-^ mice (Figures 2B, 2C, and S2A). Similarly, the frequency of CD4 T cells specific to the *Lm* protein LLO and the activation of bulk CD4 T cells were decreased in *Tlr2*^-/-^ mice (Figures S2B-S2D). Overall, these findings demonstrate that TLR2 signaling was required for the optimal priming of *Lm*-specific T cells during IV *Lm* administration, in line with the well-established role of TLR signaling in supporting T cell priming.

**Figure 2:**
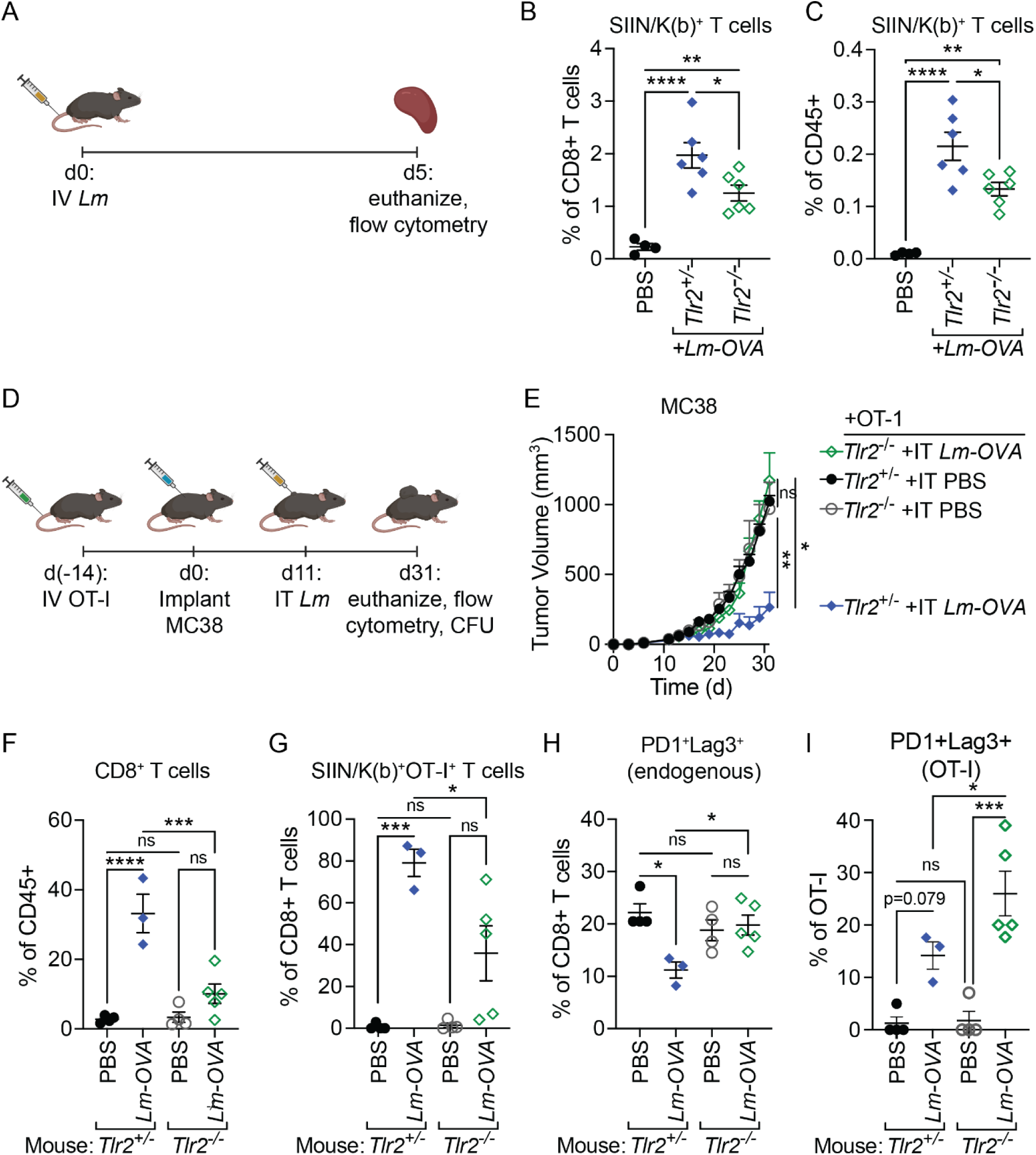
TLR2 controls the expansion and function of CD8+ T cells in IV+IT *Lm* therapy. (A) *Tlr2*^+/-^ and *Tlr2*^-/-^ littermate mice (n=6 mice/group) were infected IV with 1×10^6^ CFU of *Lm-OVA*. Splenocytes were analyzed by flow cytometry five days later. (B, C) Quantified frequencies of SIIN/K(b) tetramer-specific CD8 T cells in spleens five days after *Lm-OVA* infection presented as the frequency of CD8+ cells (B) or of CD45+ cells (C). (D) 2×10^6^ *in vitro* activated OT-I T cells were adoptively transferred to *Tlr2*^+/-^ (n=5 mice/group) or *Tlr2*^-/-^ (n=5 mice/group) littermates. Two weeks later (day 0) mice were implanted with 5×10^5^ MC38 cells and on day 11 were treated IT with 5×10^7^ CFU *Lm-OVA* or PBS. (E) Tumor growth through day 31 in mice receiving OT-I transfer and IT *Lm-OVA* or PBS. (F) Bulk CD8+ T cell frequency in tumors on day 31 (*Tlr2*^+/-^ + PBS: n=4 mice; *Tlr2*^+/-^ + IV+IT *Lm*: n=3 mice; *Tlr2*^-/-^ + PBS: n=4 mice; *Tlr2*^-/-^ + IV+IT *Lm*: n=5 mice). (G) SIIN/K(b) tetramer+ cell frequency among total CD8+ T cells in tumors on day 31. (H) Co-expression of PD1 and Lag3 on endogenous CD8 T cells on day 31. (I) Co-expression of PD1 and Lag3 on transferred OT-I T cells on day 31. Mean ± s.e.m. and one-way ANOVA with multiple comparisons (B, C, F, G, H) or two-way ANOVA (E); P: *≤0.05, **≤0.01, ***≤0.001, ****≤ 0.0001.

We then asked if TLR2 signaling played a role in supporting the CD8^+^ T cell response during IT *Lm* administration. We adoptively transferred *in vitro*-activated OT-I CD8^+^ T cells into naïve *Tlr2*^+/-^ and *Tlr2*^-/-^ littermate control mice, implanted MC38 tumors, and injected tumors IT with *Lm-OVA* eleven days later, which is sufficient to recapitulate tumor control elicited by IV+IT *Lm* (Figure 2D).^39^ Consistent with our results using IV *Lm*+IT *LmΔlgt*, *Tlr2*^-/-^ mice harboring transferred OT-I T cells failed to control their tumors in response to IT *Lm-OVA* (Figure 2E). Endpoint analysis of tumors revealed a marked reduction in the frequency of total tumor-infiltrating CD8^+^ T cells in *Tlr2*^-/-^ mice, as well as in the frequency of OT-I T cells (Figures 2F and 2G). Endogenous CD8^+^ T cells and OT-I T cells in *Tlr2*^-/-^ mice treated with *Lm-OVA* also exhibited increased expression of the inhibitory receptors Lag3 and PD1 (Figures 2H, 2I and S2E-H). Finally, control of *Lm-OVA* CFUs was significantly attenuated in tumors from *Tlr2*^-/-^ mice (Figure S2I). Together, these findings indicate that TLR2 signaling is required at the IT stage of IV+IT *Lm*, at least in part, for promoting the function of anti-*Lm* CD8^+^ T cells within tumors.

### *Lm-*triggered TLR2 signaling enhances tumor antigen uptake by DCs

To identify the immune cells that respond to *Lm* bacterial lipoproteins in tumors, we developed a fluorescent tracking approach to identify cells in tumors that: (1) express TLR2, (2) harbor *Lm*, and (3) have taken up tumor antigen. We used *Tlr2^GFP^* mice to identify TLR2-expressing cells.^59^ We injected tumors with *Lm-tagBFP*, wherein BFP expression is driven by the *actA* promoter leading to BFP fluorescence in cytosolically localized *Lm*.^60^ This approach allowed us to identify cells that harbored *Lm* and, by proxy, the cells most likely to trigger TLR2 signaling during phagocytosis of *Lm*, which can occur during the uptake of *Lm* and other TLR2 ligands.^61–64^ Finally, we implanted mCherry-expressing MC38 tumors to identify mCherry^+^ immune cells that had taken up mCherry from tumors (Figures 3A-B).

**Figure 3:**
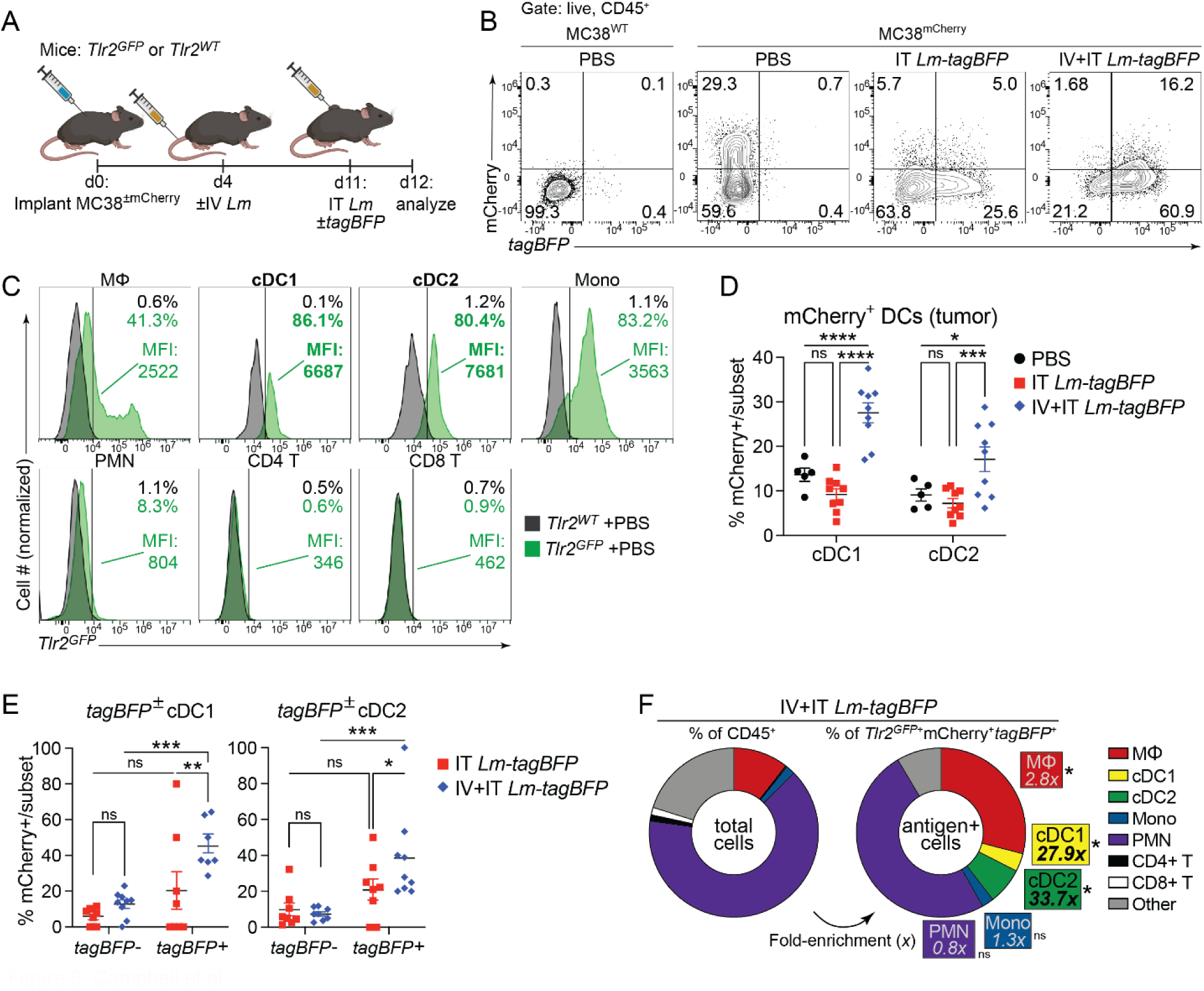
*Lm-*triggered TLR2 signaling enhances tumor antigen uptake by DCs. (A) *Tlr2^GFP^* or WT mice were implanted with 5×10^5^ MC38 cells ±mCherry (day 0) and were treated IV with 1×10^6^ CFU *Lm*±*tagBFP* or PBS (day 4) followed by 5×10^7^ CFU *Lm*±*tagBFP* (n=9 mice/group) or PBS (n=5 mice/group) IT (day 11) 24 hours prior to euthanasia and flow cytometry analysis (day 12). (B) Representative flow cytometry histograms showing mCherry and *tagBFP* fluorescence by CD45+ cells from MC38 tumors of *Tlr2^GFP^* mice treated with PBS, IT *Lm-tagBFP*, or IV+IT *Lm-tagBFP*. (C) Representative flow cytometry histograms showing GFP fluorescence by immune cell subsets in MC38 tumors of *Tlr2^GFP^* or *Tlr2^WT^* mice. (D) Frequency of mCherry+ DC subsets in MC38 tumors. (E) Cell type composition of total CD45+ versus *Tlr2^GFP^*+mCherry+*tagBFP+*CD45+ immune cells in IV+IT *Lm-tagBFP*-treated MC38 tumors showing fold-enrichment of macrophages, DC1s, and DC2s among triple-positive cells relative to their frequency among total CD45+ cells. (F) Frequency of mCherry+ among *tagBFP*± DC1s (left) and DC2s (right) in MC38 tumors of mice treated with IT *Lm-tagBFP* or IV+IT *Lm-tagBFP*. Mean ± s.e.m. and one-way ANOVA with multiple comparisons (D, E) or two-way ANOVA (F); P: *≤0.05, **≤0.01, ***≤0.001, ****≤ 0.0001.

First, we found that DCs and monocytes almost uniformly expressed TLR2 (80-100%), but that DCs expressed the highest levels of TLR2 by mean fluorescence intensity (Figures 3C and S3C-D). Lower fractions of macrophages and polymorphonuclear neutrophils (PMNs) expressed TLR2 (30% and 5%, respectively), and T cells did not express detectable levels of TLR2. We previously observed that tumor antigen uptake and presentation by DCs was enhanced in mice treated with IV+IT *Lm*, but did not identify the signals that drove this response.^39^ As DCs were highly enriched for *Tlr2^GFP^* expression in MC38 tumors, we hypothesized that the TLR2-dependent CD8^+^ T cell response following IV+IT *Lm* might be driven by increased antigen presentation by DCs. We therefore assessed the uptake of tumor antigen (mCherry) and *Lm* (*tagBFP*) by APCs within tumors. While neither IT *Lm* alone nor IV+IT *Lm* led to an increase in total mCherry^+^CD45^+^ immune cells in tumors, IV+IT *Lm* specifically increased mCherry uptake in DC1s and DC2s (Figures 3D and S3A-C). IV+IT *Lm* also led to an increase in total *tagBFP*^+^CD45^+^ immune cells (Figure 3B). This was due in part to increased *Lm-tagBFP* uptake in DC1s and DC2s with IV+IT *Lm-tagBFP* compared to IT *Lm-tagBFP* alone (Figure S3D-F). This coincided with the emergence of a substantial population of cells that took up both mCherry and *tagBFP* in IV+IT *Lm-tagBFP*-treated tumors (Figure 3B). These mCherry+*tagBFP*+ cells were also nearly all *Tlr2^GFP^*^+^ (Figure S3G), and *Tlr2^GFP^*+mCherry+*tagBFP*+ cells were significantly enriched for DC1s and DC2s (Figure 3E and S3H; 27.9x and 33.7x relative to total CD45^+^, respectively).

We next sought to determine whether increased mCherry uptake by tumor DCs was driven by the uptake of *Lm.* By stratifying tumor DCs from IV+IT *Lm-tagBFP*-treated tumors on the basis of *tagBFP* fluorescence, we observed that mCherry uptake was increased only in *Lm-tagBFP*^+^ DCs (Figure 3F). Similarly, only *Tlr2^GFP^*+ cells took up more mCherry in IV+IT *Lm*-treated tumors (Figure S3I). Intriguingly, IV+IT *Lm* significantly increased the frequency of mCherry^+^*Tlr2^GFP^*^+^ cells in the tumor-draining lymph node (tdLN) (Figure S3J). Together, these findings support our hypothesis that *Lm*-mediated TLR2 signaling supports T cell responses in tumors by modulating antigen presentation.

### IV+IT *Lm* boosts tumor antigen cross-presentation on DCs and enhances tumor-specific T cell priming

The enhancement of tumor antigen and *Lm* uptake by DC1s with IV+IT *Lm* aligned with our previous finding that tumor control with IV+IT *Lm* required the activity of tumor-specific T cells and increased tumor antigen cross-presentation by Batf3^+^ DC1s.^39^ We therefore interrogated the role of TLR2 in these aspects of the response to IV+IT *Lm* in MC38-B2m^KO^-OVA tumors. The incorporation of OVA into a MHC I-deficient tumor allows for both the detection of the OVA-derived peptide SIINFEKL bound to H-2K(b) using the 25D1.16 monoclonal antibody and transferred OT-I T cells to monitor tumor antigen-specific responses without any direct antigen presentation by tumor cells.^39,66^

We first tested the requirement for TLR2 in IV+IT *Lm*-mediated cross-presentation by treating MC38-B2m^KO^-OVA± tumor-bearing mice with IV+IT *Lm* and analyzing cross-presentation by flow cytometry 24 hr later (Figures 4A-B).^39^ As we had observed previously, CD103^+^XRC1^+^ cDC1s had the highest frequency of SIIN/K(b)^+^ in tumors, and this was further increased by IV+IT *Lm* (Figure 4C).^39^ While the fraction of SIIN/K(b)^+^ cDC2s also increased significantly with IV+IT *Lm* (Figure 4D), cDC1s remained the cell type with the highest degree of tumor antigen cross-presentation. In contrast, few to no macrophages, PMNs, or monocytes cross-presented tumor antigen (Figures S4A-S4C). Finally, increased tumor antigen cross-presentation by IV+IT *Lm* was abrogated in *Tlr2^-/-^* mice (Figures 4E-4F and S4D-S4F). Together, these findings demonstrate that IV+IT *Lm* acts via TLR2 to increase tumor antigen cross-presentation by DCs.

**Figure 4:**
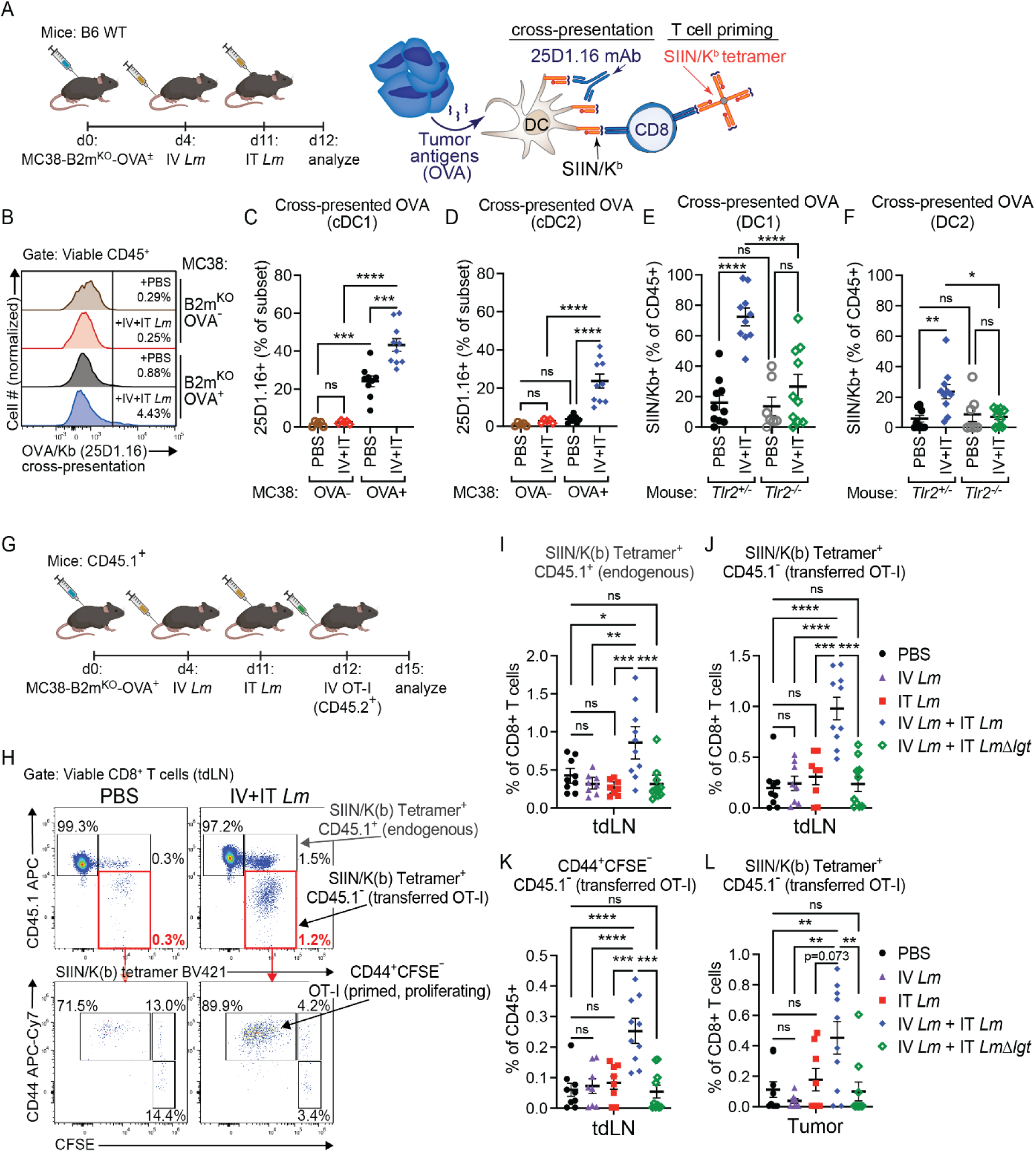
IV + IT *Lm* acts via *Lm*-specific T cells and TLR2 to induce the cross-presentation of tumor antigen and boost tumor-specific CD8 priming. (A) *Above*: WT B6 mice were implanted with 1×10^6^ MC38-B2m^KO^ cells ±OVA (day 0) and were treated with 1×10^6^ CFU *Lm* IV (day 4) followed by 5×10^7^ CFU *Lm* IT (day 11) or PBS 24 hours prior to euthanasia and analysis. *Below*: diagram depicting how surface SIIN/K(b) complexes are detected by 25D1.16 mAb staining to identify tumor antigen cross-presentation, while SIIN/K(b)-reactive TCRs are stained with SIIN/K(b) tetramer to identify tumor-specific CD8 T cells. (B) Representative flow cytometry histograms showing SIIN/K(b) staining in total CD45+ TILs in MC38. (C, D) Frequency of SIIN/K(b)+ cells among cDC1s (D) and cDC2s (E) detected by 25D1.16 staining in tumors of MC38-B2m^KO^ control (n=5 mice/group) or MC38-B2m^KO^-OVA^+^ (PBS: n=9 mice; IV+IT *Lm*: n=10 mice) tumor-bearing mice. (E, F) Frequency of SIIN/K(b)+ cells among cDC1s (E) and cDC2s (F) detected by 25D1.16 staining in tumors of MC38-B2m^KO^-OVA^+^ tumor-bearing *Tlr2*^+/-^ and *Tlr2*^-/-^ littermate mice (*Tlr2^+/-^*: n=10 mice/group; *Tlr2^-/-^* + PBS: n=8 mice; *Tlr2^-/-^* + IV+IT *Lm*: n=10 mice). (G) CD45^.1/.1^ B6 mice were implanted with 1×10^6^ MC38-B2m^KO^-OVA^+^ cells (day 0) and were treated with 1×10^6^ CFU *Lm* or PBS IV (day 4) followed by 5×10^7^ CFU *Lm*, *LmΔlgt*, or PBS IT (day 11). 24 hr later (day 12), 5×10^5^ naive, CFSE-labeled, CD45^.2/.2^ OT-Is were transferred by IV injection. After 3 days (day 15) mice were euthanized and tissues were analyzed by flow cytometry to assess OT-I infiltration and priming (PBS: n=9 mice; IV *Lm* only: n=8 mice; IT only: n=8 mice; IV+IT: n=10 mice; IV *Lm*+IT *LmΔlgt*: n=10 mice). (H) Representative flow cytometry plots demonstrating the gating strategy used to distinguish transferred OT-I (CD45.1-SIIN/K(b)+) from endogenous tumor-specific T cells (CD45.1+SIIN/K(b)+) and to assess OT-I priming and proliferation (CD44+CFSE-). (I) Frequency of endogenous SIIN/K(b)-specific CD8 T cells among CD8 T cells in the tdLN. (J) Frequency of transferred OT-I among total CD8 T cells in the tdLN. (K) Frequency of primed, proliferating OT-I among total CD45+ lymphocytes in the tdLN. (L) Frequency of transferred OT-I among CD8 T cells in MC38 tumors. Mean +/- s.e.m. and one-way ANOVA with multiple comparisons (C, D, E, F, I, J, K, L); P: *≤0.05, **≤0.01, ***≤0.001, ****≤ 0.0001.

Because IV+IT *Lm* drove TLR2-dependent cross-presentation of tumor antigens by DCs (Figure 4E-F),^39^ increased the frequency of tumor *Tlr2^GFP^*^+^mCherry^+^ cells in the tumor-draining lymph node (Figure S3J), and boosted tumor-specific CD8^+^ T cell responses following IV+IT *Lm* administration,^39^ we hypothesized that IV+IT *Lm* would lead to a TLR2-dependent increase T cell priming in lymphatic tissues. To address this question, we transferred naive, CFSE-labeled OT-I CD8^+^ T cells into congenic CD45.1^+^ MC38-B2m^KO^-OVA^+^ tumor-bearing mice after IV+IT *Lm* administration (Figure 4G). After three days, tumor, tdLN, and non tumor-draining lymph nodes (ndLN) were harvested and OT-I CD8^+^ T cells in these tissues were assessed by flow cytometry for proliferation by CFSE dilution (Figure 4H). In addition, the frequency of endogenous SIIN-specific CD8^+^ T cells was monitored (Figure 4H). IV+IT *Lm* led to a significant increase in the frequency of both endogenous tumor-specific CD8^+^ T cells and transferred OT-I T cells in the tdLN (Figures 4I, 4J, and S4G). Importantly, neither IV *Lm* alone, IT *Lm* alone, nor IV *Lm*+IT *LmΔlgt* increased the frequencies of endogenous or transferred tumor-specific CD8^+^ T cells compared to PBS treated mice (Figures 4I and 4J). The increased frequency of OT-I CD8^+^ T cells in the tdLN with IV+IT *Lm* resulted from increased OT-I proliferation as determined by increased CFSE dilution (Figure 4K), and ultimately, coincided with increased frequencies of transferred OT-I CD8^+^ T cells in the tumor (Figure 4L). Therefore, the improved tumor antigen-specific CD8^+^ T cell response elicited by IV+IT *Lm* is at least in part due to increased priming of tumor antigen-specific CD8^+^ T cells in the tdLN that is dependent on TLR2.

### TLR2 signaling decreases Foxp3 in Tregs and leads to Treg depletion in tumors

Tregs can restrict T cell priming in tumors and the tdLN by limiting tumor antigen presentation to T cells.^67–72^ Because TLR2 was required to elicit priming of tumor-specific T cells in response to IV+IT *Lm*, we hypothesized that TLR2 stimulation by IT *Lm* may modulate antigen presentation via effects on Tregs. Endpoint analysis of tumor-bearing mice treated with IV+IT *Lm* revealed a significant decrease in the frequency of tumor Tregs in WT, *Tlr5*^-/-^, and *Tlr9^-/-^*mice that did not occur in *Tlr2*^-/-^ mice (Figure 5A). This was most apparent when quantifying Tregs as a fraction of CD4^+^ T cells (Figure 5A), suggesting that the reduction in tumor Tregs was not solely the result of an influx of other cell types (e.g., PMNs or CD8^+^ T cells) but rather resulted from the specific depletion of Tregs from tumors. Combined with an increase in CD8^+^ T cells, IV+IT *Lm* led to a significantly increased ratio of CD8^+^ T cells to Tregs, which serves as a prognostic indicator of beneficial responses to immunotherapy across multiple cancer types (Figure 5B).^73,74^ Reductions in Treg frequencies and increased CD8:Treg ratios were also weakened by substituting *LmΔlgt* at the IT stage of *Lm* administration (Figures S5A-S5C). However, substituting *LmΔlgt* during IV *Lm* administration did not impact tumor Treg frequency to the same degree as substituting *LmΔlgt* during IT *Lm* administration (Figure S5C). This suggested that TLR2 signaling during IT *Lm* administration was principally responsible for reduced Treg frequencies in tumors.

**Figure 5:**
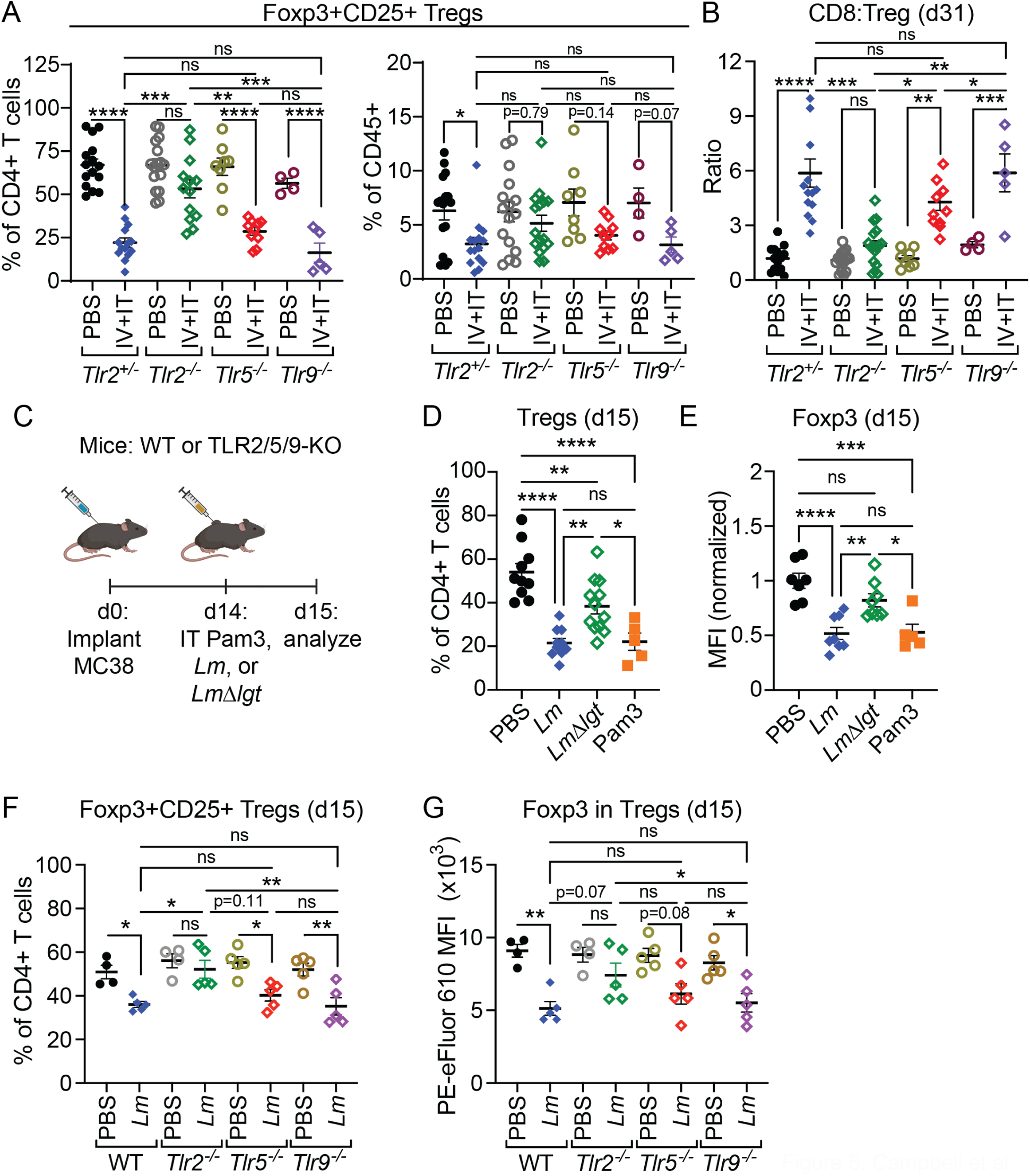
TLR2 stimulation by *Lm* mediates the depletion of Tregs from MC38 tumors. (A) Frequency of Tregs as % of CD4^+^ T cells (left) or of total CD45^+^ TILs (right) on day 31 of tumor growth in WT (*Tlr2^-/-^*; PBS: n=15 mice, IV+IT *Lm*: n=14 mice), *Tlr2^-/-^* (PBS: n=16 mice, IV+ITI *Lm*: n=13 mice), *Tlr5^-/-^* (PBS: n=8 mice, IV+IT *Lm*: n=10 mice), or *Tlr9^-/-^* (PBS: n=4 mice, IV+IT *Lm*: n=5 mice) - see Fig. 1B. (B) Ratio of CD8^+^ T cells:Tregs on day 31 of tumor growth in WT, *Tlr2^-/-^*, *Tlr5^-/-^*, or *Tlr9^-/-^* mice treated with PBS or IV+IT *Lm*. (C) WT (D-E), or WT and *Tlr2^-/-^*, *Tlr5^-/-^*, and *Tlr9^-/-^* (E-F) mice were implanted with 5×10^5^ MC38 cells (day 0) and were treated with 5×10^7^ CFU *Lm* or *LmΔlgt*, 100 ug Pam3, or PBS 24 hours prior to euthanasia and flow cytometry analysis. (D) Frequency of Tregs as % of CD4^+^ T cells 24 hr post-IT PBS (n=10 mice), *Lm* (n=9 mice), *LmΔlgt* (n=12 mice), or Pam3 (n=5 mice). (E) Foxp3 fluorescence within Foxp3^+^CD25^+^ tumor Tregs 24 hr post-IT. (F) Frequency of Tregs as % of CD4^+^ T cells in WT (PBS: n=4 mice, *Lm*: n=5 mice), *Tlr2^-/^* (PBS: n=4 mice, *Lm*: n=5 mice)*^-^*, *Tlr5^-/-^* (n=5 mice/group), and *Tlr9^-/-^*(n=5 mice/group) 24 hr post-IT. (G) Foxp3 fluorescence within Foxp3^+^CD25^+^ tumor Tregs 24 hr post-IT PBS or *Lm*. Mean +/- s.e.m. and one-way ANOVA with multiple comparisons (A, B, D, E, F, G); P: *≤0.05, **≤0.01, ***≤0.001, ****≤ 0.0001.

Increased tumor antigen cross-presentation and trafficking to the tdLN occurred within 24 hr of treatment with IV+IT *Lm* (Figures 4B-F and S3J), leading us to hypothesize that the impact of TLR2 on tumor Tregs would be similarly rapid. To investigate the immediate impact of TLR2 signaling on tumor Tregs, we treated MC38 tumor-bearing mice with *Lm*, *LmΔlgt*, or Pam3 and analyzed the tumors by flow cytometry 24 hr later (Figure 5C). IT *Lm* led to a marked reduction in the frequency of tumor-infiltrating Tregs (Figure 5D). This was accompanied by a significant reduction in the MFI of Foxp3 within remaining Foxp3^+^ Tregs, which is often associated with reduced Treg suppressive function (Figure 5E).^75^ These changes were confined to the tumor, as the frequency of Tregs was not reduced by IT *Lm* in proximal (tdLN) or distal (ndLN and spleen) lymphatic tissues (Figure S5D). While IT Pam3 was sufficient to phenocopy Treg frequency reduction and Foxp3 downregulation observed with IT *Lm*), these effects were significantly muted with IT *LmΔlgt* (Figures 5D-E and S5E). Similarly, *Tlr2^-/-^* but not *Tlr5^-/-^ or Tlr9^-/-^* mice failed to exhibit reduced tumor Treg frequency or Foxp3 expression following IT *Lm* administration (Figures 5E-G). These results indicate that TLR2 signaling in the TME is both necessary and sufficient to reduce Foxp3 expression and Treg frequencies in response to IT *Lm*, which may underlie the dependence of IV+IT *Lm* on TLR2 but not TLR5 or TLR9.

To determine whether *Lm*-mediated TLR2 stimulation led to increased generation of “ex-Tregs”, Tregs that lose their suppressive function due to transcriptional reprogramming upon loss of Foxp3.^76,77^ To test this possibility, we used Foxp3 lineage-trace mice in which the *Foxp3* promoter drives the expression of a GFP-Cre fusion protein, which labels Foxp3^+^ cells with GFP and excises a *lox-stop-lox* cassette upstream of an IRES-RFP element at the constitutively expressed *Rosa26* locus (Figure 6A).^78,79^ IT Pam3 led to decreased MFI for GFP and Foxp3 antibody staining, which corresponded to an increase in the formation of GFP^-^RFP^+^ ex-Tregs in tumors (Figures 6B-D). These findings confirm that TLR2 signaling in tumors disrupts Foxp3 expression and lineage stability in Tregs.

**Figure 6:**
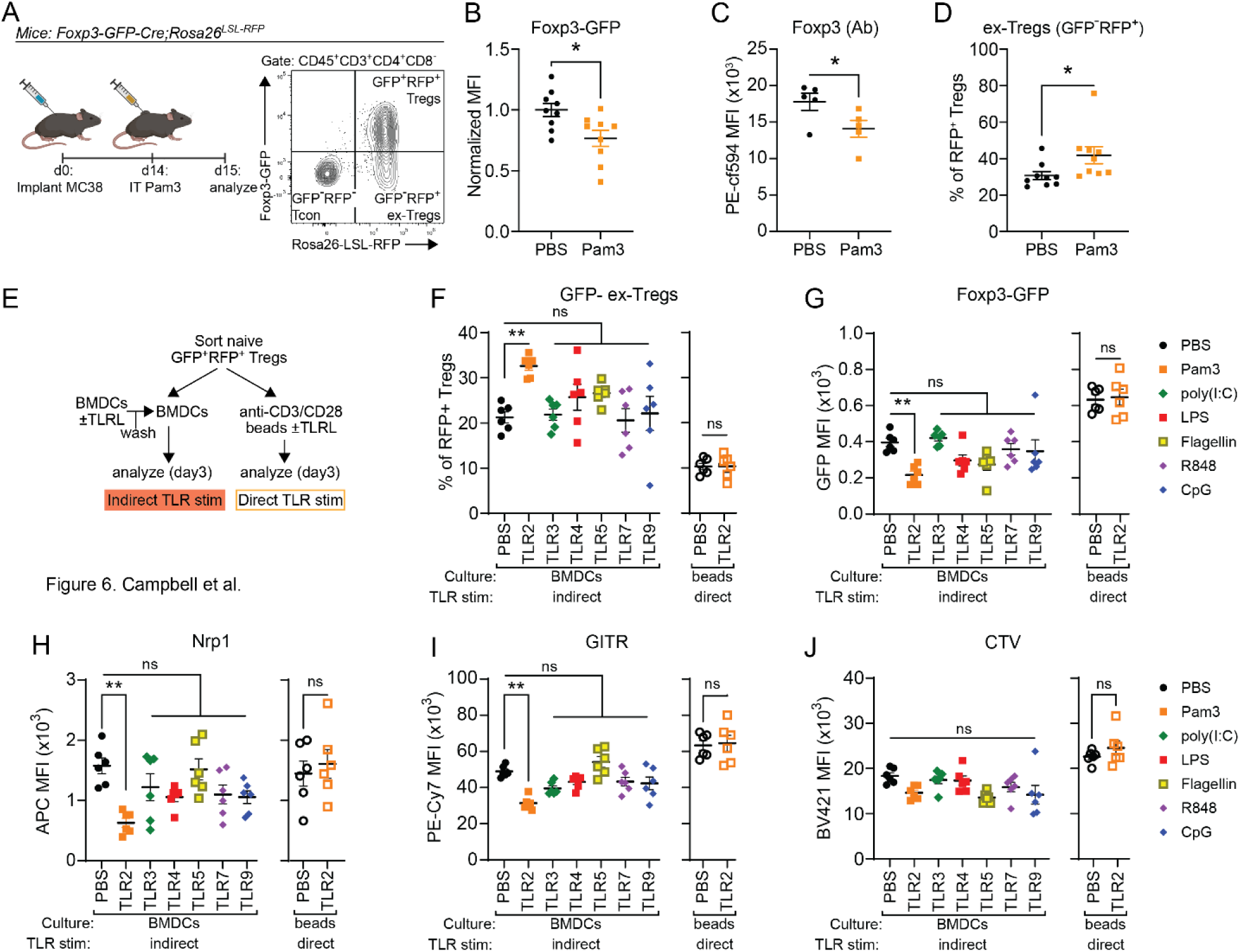
TLR2 signaling destabilizes Tregs via a cell-extrinsic mechanism. (A) *left*: Foxp3 lineage-trace mice were implanted with 5×10^5^ MC38 cells (day 0) and were treated IT with 100 ug Pam3 or PBS (day 14) 24 hours prior to euthanasia and flow cytometry analysis of cells in MC38 tumors. *Right*: representative flow cytometry plot showing CD4^+^ T cells in MC38 tumors of Treg lineage trace mice with conventional CD4^+^ T cells (Tcon), stable Tregs, and ex-Tregs labeled. (B) MFI of *Foxp3^GFP-hCre^* expression within Foxp3^+^CD25^+^ Tregs (n=10 mice/group). (C) MFI of Foxp3 antibody stain within Foxp3^+^CD25^+^ Tregs (n=5 mice/group). (D) Frequency of GFP^-^RFP^+^ ex-Tregs among total RFP+ cells. (E) Experimental outline for Treg:BMDC co-culture to test the ability of TLRL to destabilize Tregs *ex vivo* via direct (Treg-intrinsic) or indirect (Treg-extrinsic) TLR signaling (n=2 mice, n=3 technical replicates/ea). (F) Frequency of GFP^-^RFP^+^ ex-Tregs among total RFP^+^ Tregs cultured with TLR-stimulated or unstimulated BMDCs (left) or ɑCD3/CD28 beads with or without Pam3 (right) after 72 hr co-culture. (G, H, I, J) MFI of *Foxp3^GFP-hCre^* (G), Nrp1 (H), GITR (I), and CTV (J) within RFP^+^ Tregs after 72 hr co-culture. Mean +/- s.e.m. and Student’s t-test (B, C, D) or one-way ANOVA with multiple comparisons (F, G, H, I); P: *≤0.05, **≤0.01, ***≤0.001, ****≤ 0.0001.

### TLR2 signaling in DCs uniquely destabilizes Treg Foxp3 expression

The dependence of *Lm*-mediated Treg destabilization in tumors upon TLR2, but not TLR5 or TLR9, led us to ask whether TLR2 exhibits a unique capacity to destabilize Treg expression of Foxp3. Multiple reports using *in vitro* and *in vivo* systems have identified various Treg-intrinsic effects of TLR2 stimulation,^102–106^ leading us to use a reductionist approach in which sorted, CTV-labeled naive Tregs were co-cultured *in vitro* with TLR-stimulated bone marrow-derived dendritic cells (BMDCs) or stimulated alone with anti- CD3/anti-CD28-coated microbeads with or without TLR ligands (Figure 6E). Consistent with our findings *in vivo*, Tregs cultured with TLR2-stimulated BMDCs downregulated GFP and converted to ex-Tregs to a significantly greater extent than Tregs cultured with unstimulated BMDCs (Figures 6F and 6G). Tregs cultured with TLR2-stimulated BMDCs also uniquely exhibited reduced expression of Nrp1 and GITR, two markers of functionally suppressive Tregs whose loss is associated with Treg lineage destabilization (Figures 6H and 6I).^80,81^ Notably, these effects were specific to TLR2, as BMDCs stimulated by any other TLR failed to cause Tregs to downregulate Foxp3, Nrp1, or GITR (Figures 6F-I). In contrast, direct TLR2 stimulation of Tregs had no impact on Treg phenotype (Figures 6F-6J). Importantly, Treg proliferation (quantified by CTV dilution) did not differ between these conditions, ruling out the possibility that TLR2-mediated changes in Treg phenotype were due to differences in Treg activation (Figure 6J). Finally, to confirm that cell-intrinsic TLR2 signaling in Tregs did not mediate Foxp3 destabilization *in vivo*, we treated tumor-bearing mixed bone marrow chimeric mice reconstituted with congenically marked *Tlr2*^+/-^ and *Tlr2*^-/-^ cells with IT Pam3 and found that reduced Treg frequency and Foxp3 expression were independent of Treg genotype (Figures S5F-S5H). Altogether, our results from both *in vivo* and *in vitro* systems establish that *Lm*-mediated TLR2 signaling in APCs, but not directly in Tregs, uniquely drives tumor Treg destabilization in response to IT *Lm* administration.

### TLR2-mediated reduction of tumor Tregs is required for the efficacy of IV+IT *Lm* therapy

The unique impact of TLR2 stimulation on Treg lineage stability *in vivo* and *in vitro* led us to hypothesize reducing tumor Treg frequency was part of the mechanism by which TLR2 signaling exerted its unique effect on tumor control during IT *Lm* administration in IV+IT *Lm*-treated mice. To test this model, MC38 tumors were implanted into *Foxp3^DTR^* mice to enable conditional depletion of Foxp3^+^ Tregs by DT administration, independently of TLR2 signaling.^82^ Next, tumor-bearing mice received IV *Lm* followed by one of four IT injections: 1) IT *Lm* to deplete Tregs via TLR2 and elicit tumor control (positive control); 2) IT *LmΔlgt*, which fails to deplete Tregs and does not elicit tumor control (negative control); 3) a single low dose of IT DT to transiently deplete tumor Tregs independently of TLR2 and *Lm* antigens, or 4) IT *LmΔlgt* plus a single low dose of DT to transiently deplete tumor Tregs independently of TLR2 and provide *Lm* antigens to reactivate *Lm*-specific CD8^+^ T cells (Figure 7A). Strikingly, only mice treated with IT *LmΔlgt*+DT controlled tumors comparably to mice receiving IT *Lm* (Figure 7B). This confirmed that TLR2-mediated Treg depletion by IT *Lm* was required for tumor control, establishing a mechanistic basis for the specific requirement for TLR2 signaling during IT *Lm* administration to elicit tumor control in response to IV+IT *Lm.* In addition to tumor control, the reduced Tregs and increased CD8^+^ T cell frequencies and increased CD8^+^ T cell:Treg ratio observed with IT *Lm* were partially restored with IT *LmΔlgt+*DT (Figures S6A-6C). However, mice treated with IT *LmΔlgt*+DT did not control *Lm* CFUs in tumors to the same extent as IV+IT *Lm*-treated mice, suggesting that TLR2 signaling contributes to *Lm* control in tumors via a mechanism that is distinct from but may also require Treg depletion (Figure S6D). Notably, the failure of IT DT alone to slow tumor growth demonstrates that Treg depletion by TLR2 is necessary but not sufficient for the activity of IV+IT *Lm*, analogous to the failure of IV *Lm*+IT Pam3 to fully control tumors (Figure 1H). This implies that both Treg destabilization by TLR2 and the reactivation of *Lm*-specific CD8^+^ T cells in tumors by *Lm* antigens during IT *Lm* administration are required for tumor control.

**Figure 7:**
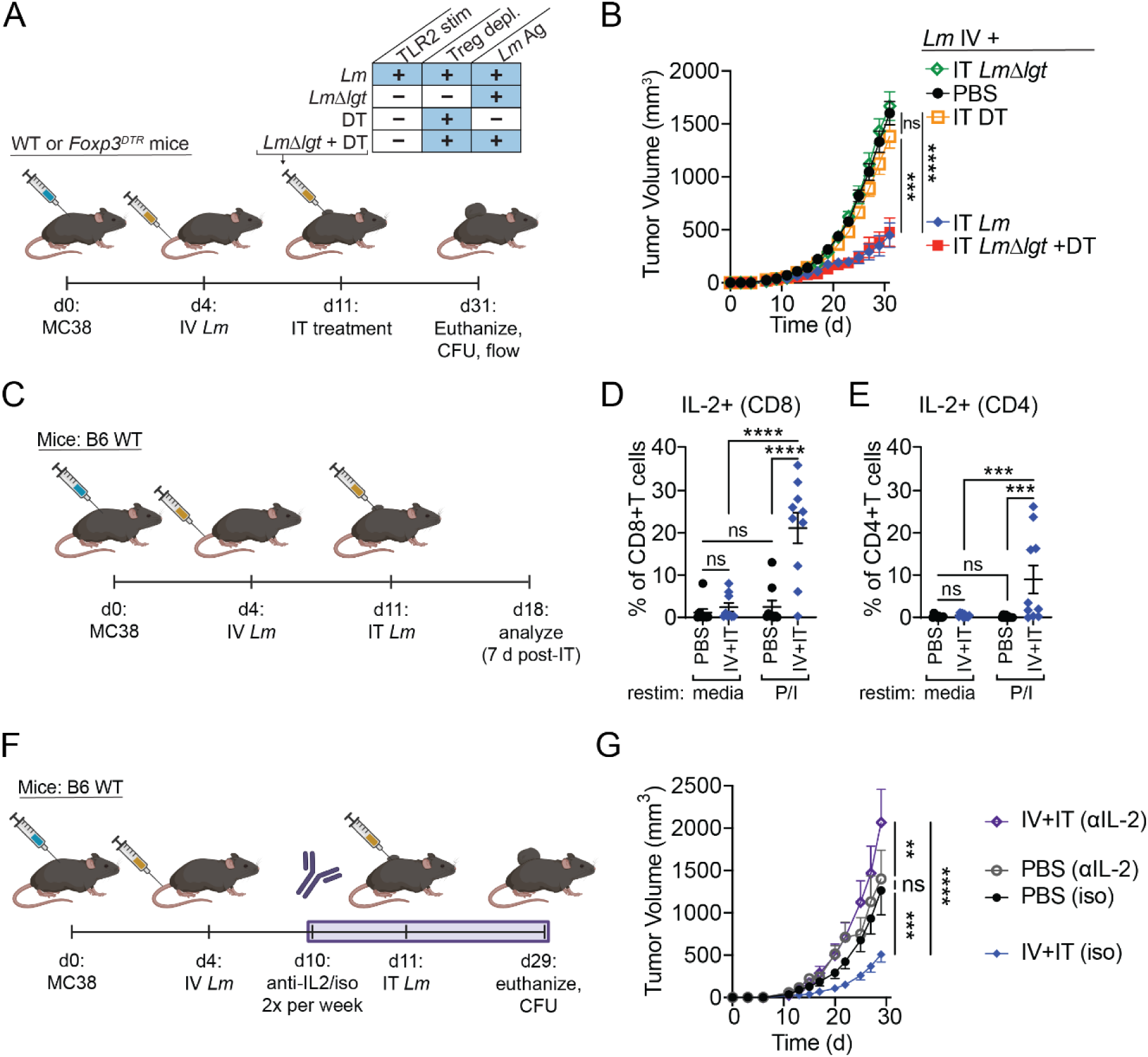
Treg depletion by TLR2 is essential for IV+IT *Lm* efficacy. (A) Experimental outline for determining whether Treg depletion by IT *Lm* is required for tumor control in response to IV+IT *Lm* (n=10 mice/group). (B) MC38 tumor growth following treatments indicated in A. (C) Experimental outline for testing if IV+IT *Lm* boosts IL-2 production in tumor T cells (n=10 mice/group). (D, E) IL-2 production in tumor-infiltrating CD8 T cells (G) and CD4 T cells (H) following 6 hr restimulation with media or PMA+ionomycin. (F) Experimental outline for determining the necessity of IL-2 for tumor control in IV+IT *Lm* (PBS + isotype: n=9 mice; PBS + αIL-2: n=10 mice; IV+IT *Lm* + isotype: n=11 mice; IV+IT *Lm* + αIL-2: n=9 mice). (G) MC38 tumor growth in the presence of an IL-2 depleting antibody or isotype control. Mean +/- s.e.m. and two-way ANOVA (B, G) or one-way ANOVA with multiple comparisons (D, E); P: *≤0.05, **≤0.01, ***≤0.001, ****≤ 0.0001.

### IV+IT *Lm* enhances IL-2 signaling in tumor-infiltrating CD8^+^ T cells to mediate tumor control

We previously observed that tumor-specific CD8^+^ T cells exhibited reduced PD1 and Lag3 expression during endpoint analysis of tumors treated with IV+IT *Lm*.^39^ While this is consistent with TLR2 driving *de novo* tumor-specific CD8^+^ T cell priming (Figures 4H-L), we hypothesized that the TLR2-mediated reduction in Treg-mediated suppression and increased tumor antigen cross-presentation would act synergistically to reduce exhaustion in preexisting tumor-specific CD8^+^ T cells. We therefore measured PD-1 and Lag3 expression on tumor-specific T cells 24 hours after IT *Lm* injection in MC38-B2m^KO^-OVA^±^ tumor-bearing mice (Figures 4A and S7A). This revealed that IV+IT *Lm* rapidly reduced the expression of both PD1 and Lag3 on tumor-specific CD8^+^ T cells (Figures S7B-S7E). In addition, expression of the high-affinity IL-2Rα chain (CD25) was increased on both tumor-specific and bulk CD8^+^ T cells with IV+IT *Lm* (Figures S7F-G), a phenotype associated with increased antitumor cytotoxic T cell activity.^83^ Together, these changes in tumor-specific CD8^+^ T cell phenotypes suggest that IV+IT *Lm* rapidly reprograms preexisting tumor-specific CD8^+^ T cells in tumors in addition to promoting *de novo* priming.

CD25 is upregulated by TCR signaling along with its ligand IL-2, which together create an autocrine signaling loop that supports T cell proliferation and effector function.^84^ Treg competition for IL-2 can limit IL-2 signaling to CD8^+^ T cells, leading to reduced CD25 expression and reduced.^84^ Therefore, we hypothesized that IV+IT *Lm* may boost IL-2 production in tumor-infiltrating CD8^+^ T cells by reducing tumor Treg frequency and increasing antigen presentation to CD8^+^ T cells. Indeed, IV+IT *Lm* significantly increased IL-2 production by tumor CD4^+^ and CD8^+^ T cells (Figures 7C-7E and S7H). Furthermore, anti-IL-2 antibody blockade beginning one day before IT *Lm* completely abrogated tumor control with IV+IT *Lm* and trended towards increasing the number of *Lm* CFUs recovered from tumors (Figures 7I, 7J, and S7I). Altogether, these results indicate that IL-2 signaling is essential for eliciting tumor control with IV+IT *Lm*. Mechanistically, these results are consistent with *Lm*-mediated TLR2 signaling reducing tumor Treg frequency and increasing tumor antigen cross-presentation, which together increase IL-2 production and signaling by tumor-specific CD8^+^ T cells to enhance their capacity to control tumors.

## Discussion

Although TLRs are widely appreciated to enhance adaptive immune responses, the mechanisms by which they do so, and the specific effects of signaling by different TLRs on adaptive immunity within the TME, remain incompletely understood. Here, using an IV+IT *Lm* regimen to elicit a potent CD8^+^ T cell-dependent anti-tumor response, we uncovered an essential and multifactorial role for TLR2 signaling at both the IV and IT phases of *Lm* administration (Figure 8). During IV *Lm* administration, TLR2 contributed to the priming of *Lm*-specific CD8^+^ T cells, in line with the well-characterized role of TLRs in promoting the priming of pathogen-specific T cells (Figure S8A). However, to our surprise, TLR2 was also critical during IT *Lm* administration. This was due to TLR2 signaling uniquely driving the destabilization of Foxp3 expression in tumor Tregs during IT Lm administration, leading to reduced Treg frequency and immunosuppression in tumors (Figure S8B). Treg depletion enhanced the frequency and functionality of *Lm*-specific CD8^+^ T cells within tumors and reversed CD8^+^ T cell exhaustion. These effects synergized with direct TLR2 signaling to enhance the uptake of both bacteria and tumor antigens and increased cross-presentation of tumor antigens on conventional DCs, leading to increased priming of tumor-specific CD8^+^ T cells in the tdLN, enhanced CD8^+^ T cell recruitment to tumors, and decreased expression of PD-1 and Lag-3 on CD8^+^ T cells in tumors (Figure S8C). These effects were due, at least in part, to increased IL-2 production and signaling in tumor CD8^+^ T cells, likely also as a result of reduced tumor Treg competition for IL-2 (Figure S8C). Thus, TLR2 signaling has the unique ability to promote CD8^+^ T cell-dependent antitumor immunity by destabilizing Tregs within tumors, revealing a novel adjuvant function of TLR2 in the context of cancer immunotherapy.

The requirement for TLR2 signaling by *Lm* to mediate tumor control with IV+IT *Lm* provides clear experimental evidence demonstrating that TLR stimulation by *L. monocytogenes* contributes to its antitumor activity.^7,85^ Until now, the importance of TLR signaling for bacterial cancer therapy was most strongly supported by the clinical success of BCG, which also strongly activates TLR2 but appears to require the activation of multiple TLRs to elicit tumor control.^86,87^ Interestingly, the only other study examining *L. monocytogenes*-mediated TLR2 signaling in the setting of cancer found that TLR2 signaling in tumor cells, and not immune cells in the TME, impacted tumor progression.^88^ In line with our results, another study found that *Akkermansia*-mediated TLR2 stimulation increased the efficacy of IL-2 cancer therapy.^89^ A variety of small-molecule TLR2 agonists have also been shown to inhibit tumor growth in preclinical mouse cancer models, bolstering interest in the anti-cancer efficacy of TLR2-activating drugs.^90–92^ Our results expand upon these findings by demonstrating that TLR2 signaling mediated by *Lm* lipoproteins provides immune-mediated control of tumors in the absence of additional therapy such as IL-2. Together, results from BCG, *Akkermansia*, and *Lm* indicate that bacterial TLR2 ligands can mediate antitumor immunity, and make a compelling case to broaden the direction of clinical investigations from the activation of endosomal TLRs to surface TLRs as adjuvants for antitumor immunity.^15,17^

TLR2 signaling is known to be important for combating primary *L. monocytogenes* infection in the liver and the spleen, but its role in CD8^+^ T cell responses to secondary infection is inconclusive, leading to the consensus that TLR2 signaling exerts its protective effects solely through innate immunity.^7,25–31^ In the broader context of antibacterial adaptive immunity, it has often been argued that TLR signaling is minimally important, and can be bypassed, for the establishment of adaptive immune responses because patients with TLR deficiencies typically have reduced susceptibility to bacterial infection over time.^32,33^ However, here we found that TLR2 was required during both IV (primary) and IT (secondary) *Lm* administration to support CD8^+^ T cell-mediated tumor control and *Lm* bacterial burden within tumors. The requirement for TLR2 signaling to mediate control of secondary *Lm* infection in our study, but not in previous reports, likely stems from our studying *Lm* in the context of cancer, where *Lm* can colonize tumors and interact with a unique set of immune cells recruited to and shaped by the TME.^39^ The TME is highly immunosuppressive, which may render individual anti*-Lm* CD8^+^ T cells less effective.^93,94^ Similarly, the frequency of *Lm*-specific CD8^+^ T cells in tumor tissue may be limited by the scarcity of nutrients or signals that promote proliferation and survival, such as IL-2.^95^ In either case, the failure of adaptive immunity to effectively control *Lm* in the absence of TLR2 signaling during IT *Lm* administration demonstrates more generally that innate immune signals not only support the priming of adaptive immune cells, but also support their function upon reactivation in response to secondary infection in immunosuppressive environments. Therefore, the immune response to *Lm* in tumors highlights mechanisms by which TLR signaling can contribute at both early and late stages of adaptive immune responses.

TLR2-mediated destabilization and depletion of Tregs in the TME critically contributed to tumor control by IV+IT *Lm*. This likely underlies the specific requirement for TLR2 during IT *Lm* administration, while other TLRs are dispensable, and may explain the requirement for TLR2 in adaptive anti*-Lm* immune responses in tumors but not in other tissues. The ability of IT *Lm* to deplete Tregs is consistent with a recent study that showed that systemic *Lm* infection disrupted Treg suppression and reduced Treg frequency in the spleen.^96^ However, the authors did not identify the signal driving these phenotypes, which we have identified here was likely TLR2 signaling. Indeed, others have shown that TLR2 signaling by small-molecule ligands negatively regulates Treg function, but did not test whether TLR2 stimulation by live bacteria or bacterial products have the same result.^97–101^ In contrast, we found no evidence of prior reports that tied TLR5 stimulation by *Lm* to Treg frequency or function, while the sole study to investigate a link between TLR9, Tregs, and *Lm* found that TLR9 signaling supported the generation of peripherally-induced Tregs from naive CD4^+^ T cells.^102^ Our work therefore bridges a gap between these findings by demonstrating that bacterial TLR2 ligands can disrupt Treg suppression in the TME, an effect that is specific to TLR2 and no other TLRs. However, it is noteworthy that Treg depletion did not lead to reduced bacterial burden in our system, suggesting that TLR2 contributes to anti-*Lm* immunity by multiple mechanisms, and not solely by negatively regulating Tregs.^61,103^

Prior reports which found that TLR2 mediated the disruption of Treg suppression proposed that this effect was driven by T cell-intrinsic TLR2 signaling within Tregs or effector T cells.^97,104^ In contrast, we did not observe evidence of TLR2 expression in Tregs, CD4^+^ conventional T cells, or CD8^+^ T cells, nor did we find evidence that T cell-intrinsic TLR2 signaling contributed to tumor control. This suggests that the effect of TLR2 on both Tregs and CD8^+^ T cells in tumors is mediated by TLR2 signaling in another cell type in the TME, which is likely DCs given their uniformly high expression of *Tlr2^GFP^*and strong capacity to influence T cell responses. In particular, our observation that DC1s cross-present tumor antigens and take up *Lm* in response to IV+IT *Lm*, and our previous finding that *Batf3* expression is required for IV+IT *Lm* therapy,^39^ implies that DC1s mediate the effects of TLR2 signaling on T cells with IV+IT *Lm*. Our findings therefore underscore the diversity of methods by which TLR2 signaling can regulate T cell immunity,^8^ in this case via a T cell-extrinsic mechanism that is translated by other immune cells into an instructive cue.

TLR2 signaling promoted CD8^+^ T cell responses by enhancing tumor antigen uptake and cross-presentation by DCs. This finding is consistent with previous reports that small-molecule TLR2 ligands covalently linked to tumor antigens can enhance cross-presentation to boost antitumor immunity.^105–109^ Similarly, the activation of tumor-nonspecific bystander CD8^+^ T cells in the TME can provide immunostimulatory signals to APCs that boost cross-presentation of tumor antigens and subsequent epitope spreading.^110,111^ Our results suggest that IV+IT *Lm* operates by a combination of these mechanisms and Treg depletion because both IV *Lm* and IT *Lm* were necessary but insufficient to increase cross-presentation and tumor-specific T cell priming. In our combined model, IV *Lm* potentiates cross-presentation by generating *Lm*-specific T cells which provide signals such as IFN-γ that promote cross-priming by DCs, as has previously been observed with other therapies that activate tumor-nonspecific bystander CD8^+^ T cells.^110–113^ In turn, IT *Lm* provides both bacterial TLR2 ligands (to drive DC-intrinsic TLR2 signals and deplete tumor Tregs) and *Lm* antigens (to reactivate *Lm*-specific CD8^+^ T cells generated by IV *Lm*) which synergize to enhance tumor antigen uptake and cross-presentation by DCs. This model is consistent with previous work from our lab which showed that IFN-γ is strongly induced in CD8^+^ T cells by IV+IT *Lm* and that IFN-γ produced by *Lm*-specific CD8^+^ T cells is required for tumor control.^39^ Furthermore, we have shown that blockade of IFN-γ or depletion of CD8^+^ T cells abrogated enhanced cross-presentation mediated by the depletion of CXCR3^+^ Tregs from MC38 tumors.^114^ This model is particularly intriguing because it would reveal a novel mechanism by which the innate and adaptive immune response against *Lm* within tumor tissue can together mediate tumor antigen cross-presentation and enhance antitumor immunity without additional immunotherapeutic treatments, the co-expression of tumor antigens by *Lm*, or the direct infection of tumor cells by *Lm*.^4,7,115^

In summary, our findings have important implications for the development of immunotherapies that act on innate immune cells to promote the T cell response, including bacterial immunotherapies. First, tumor-specific CD8+ T cell immunity can be increased in the absence of genetically engineering the expression of tumor antigens into bacteria, instead relying upon the ability of bacteria-specific CD8+ T cells and bacterial TLR ligands to synergistically mediate tumor antigen cross-presentation. This finding has tremendous potential in light of many recent discoveries that several types of cancer may naturally harbor numerous microbes.^92,118,119^ Our findings suggest that eliciting T cell responses against such microbes within tumors can cause epitope spreading to tumor antigens, yielding therapeutically efficacious antitumor immunity. Second, TLR2 signaling increases antigen uptake by APCs and inhibits Treg suppression, raising the importance of targeting TLR2 therapeutically. While several TLR agonists have been tested in clinical trials, TLR2 has not been the focus of these efforts.^10,13–16^ However, our results, along with several other recent studies,^42,120–125^ indicate that TLR2 stimulation may be a more potent agonist of T cell-mediated antitumor immunity than formerly realized, likely due to its unique inhibition of Treg-mediated immunosuppression within tumors. Thus, our work supports the development of cancer immunotherapies that stimulate TLR2. Third, enhanced tumor antigen presentation in tdLN can enhance the systemic immune response against tumor antigens, providing a mechanism for intratumoral immunotherapies to ignite systemic adaptive immune responses that could be effective against disseminated metastatic cancers. In conclusion, by characterizing the molecular and cellular mechanisms underlying the TLR2 dependency of IV+IT *Lm* therapy, we highlight how this receptor can be leveraged to elicit improved antitumor immunity.

## Limitations of the study

All experiments were conducted in preclinical mouse models of disease, which can fail to recapitulate responses to therapies in humans. Additional studies using humanized mice may provide insight into the translatability of the specific mechanisms identified in this work to the human immune system. The primary strain used in this study, *Listeria monocytogenes ΔactA*, has a tolerable safety profile in human clinical trials, but has been observed to form biofilms on medical implants in some patients during clinical trials. Mutations that reduce biofilm formation and the extracellular viability of *L. monocytogenes* may be considered for clinical development based upon the principles outlined in this work.

## Resource Availability

### Lead Contact

Further information and requests for resources and reagents should be directed to and will be fulfilled by the lead contact, Michel DuPage (dupage{at}berkeley.edu).

### Materials availability

*Listeria monocytogenes* strains used in the study will be made available from the lead contact upon request.

### Data and code availability

Any additional information required to reanalyze the data reported in this paper is available from the lead contact upon request.

## Acknowledgements

We thank Prof. Greg Barton (UC Berkeley) for the provision of *Tlr2^-/-^*and *Tlr2^GFP^* mice. We thank Prof. Jeff Bluestone (UCSF) for the provision of Foxp3 lineage trace mice and MC38 and MC38-OVA cells. We thank the UC Berkeley Cancer Research Laboratory Flow Cytometry Core staff for providing technical assistance and training in flow cytometry and FACS. We thank Profs. David Raulet, Ellen Robey, Russell Vance, and Greg Barton (UC Berkeley) for thoughtful discussion and for providing feedback on this manuscript. We thank Antoine Sacquet, Elina Wells, Jacob Williams, and others for their assistance in performing experiments. This research was supported by by 1DP2CA247830-01, 1R01AI27655, and 1P01AI063302 from the National Institutes of Health (NCI) and C23CR5612 from the UC CRCC Cancer Research Coordinating Committee. T.C. is supported by the UC CRCC Fellowship. J.G.C is a HHMI Gilliam Fellow. M.D. is a Pew-Stewart Scholar and a St. Baldrick’s Scholar with generous support from Hope with Hazel.

## Author Contributions

Conceptualization and methodology, M.D., T.C, and D.A.P..; investigation, T.C., J.G.C., S.F., D.G.V., S.H., J.V., N.F.H., and J.Y.; writing – original draft, M.D. and T.C.; writing – review & editing, M.D., T.C., and D.A.P..; supervision, M.D., T.C., and D.A.P.; funding acquisition, M.D. and D.A.P.

## Methods

### Experimental models and subject details

#### Animal studies

C57BL/6J wild-type (JAX:000664) and CD45.1+ (JAX:002014) mice were obtained from Jackson laboratories and bred in house. OT-1 transgenic mice were obtained from Taconic (Catalog#: 2334) and bred in house. *Tlr2^-/-^* (JAX:004650) and *Tlr2^GFP^* (JAX:031822) mice were a gift from the Barton lab at the University of California, Berkeley. Foxp3 lineage trace mice were generated by crossing *Foxp3^eGFP-hCre^* mice (JAX:023161) with *Rosa26^LSL-RFP^*mice^126^ and were a gift from the Bluestone lab at the University of California, San Francisco. For tumor studies, syngeneic C57BL/6J mice of the above genotypes were inoculated with 5.0×10^5^ MC38, 5.0×10^5^ MC38^mCherry^, 1.0×10^6^ MC38-B2m^KO^, 1×10^6^ MC38-B2m^KO^-OVA, or 2.0×10^5^ B16F10 tumor cells in PBS subcutaneously on the right flank. For KP sarcoma experiments, 2.0×10^5^ KP sarcoma cells were implanted intramuscularly in the right hind leg.^39^ Tumor measurements were performed blindly across the entire experiment by a single operator measuring three dimensions of the tumor with calipers every second day beginning when tumors were palpable. All experiments were conducted according to the Institutional Animal Care and Use Committee guidelines of the University of California, Berkeley.

#### Cell lines

MC38 and MC38-OVA cell lines were kindly provided by Dr. Jeff Bluestone’s lab. The MC38-β2m^KO^ and MC38-β2m^KO^-OVA cell lines were generated by our lab by CRISPR-Cas9-mediated knockout of *B2m* followed by FACS to purify H2-K(b)^-^H2-D(b)^-^ cells using a FACSAria™Fusion cell sorter (BD).^127^ The MC38^mCherry^ cell line was generated by our lab as detailed below. All cell lines were maintained in DMEM (GIBCO) supplemented with 10% FBS, sodium pyruvate (GIBCO), L-Glutamine (GIBCO), and penicillin-streptomycin (GIBCO). Tumor cells were grown at 37°C with 5% CO2.

### Method Details

#### Generation of mCherry-expressing MC38

The MC38^mCherry^ cell line was generated by transduction of MC38 cells with an mCherry-encoding lentiviral construct as described previously.^128^ Briefly, VSV-G, psPax2, and mCherry-encoding vector targeting plasmid were transfected into LentiX-293T cells (Takara Bio) at a molecular ratio of 1:3:4 using Lipofectamine™ Transfection Reagent (Thermo Fisher) on day 0. The following day (day 1), LentiX-293T media was swapped for fresh media. On day 4, lentiviral media was collected and filtered through a 0.45 um syringe-driven filter. MC38 cells were suspended in lentiviral media in the presence of 4 ug/mL polybrene (Sigma-Aldrich) and incubated at 37°C with rotation for 1 hr. Following incubation, cells were resuspended in fresh media and expanded through serial passaging. mCherry^+^ MC38 cells were isolated to >95% purity by multiple rounds of FACS using a FACSAria™Fusion cell sorter (BD).

#### L.monocytogenes strains

All strains of *L. monocytogenes* were derived from the wild-type 10403S strain. The *Lm* constructs were based on an attenuating deletion of the actA gene (Δ*actA*). *LmΔlgt* was generated by introduction of an in-frame deletion of *lgt*.^129^ *Lm-OVA* expresses an LLO-OVA fusion encompassing amino acids 1-441 of Listeriolysin O (LLO) and OVA cloned in frame of LLO1-441 under the control of the *hly* promoter in the pPL2 integration vector.^130,131^ *Lm-TagBFP* expresses a secreted TagBFP protein under the control of the *actA* promoter using the pPL2 integration vector.^60,131^ All strains were cultured in filter-sterilized nutrient-rich Brain Heart Infusion (BHI) media (BD Biosciences) containing 200 μg/mL streptomycin (Sigma-Aldrich).

#### Intravenous and Intratumoral Listeria infection

Overnight cultures were grown in BHI + 200 μg/mL streptomycin at 30°C. The following day, bacteria were grown to logarithmic phase by diluting the overnight culture in fresh BHI + 200 μg/mL streptomycin and culturing at 37°C shaking. Log-phase bacteria were washed and frozen in 9% glycerol/PBS. For intravenous infections, frozen stocks were diluted in PBS to infect via the tail vein with 1×10^6^ CFU log-phase bacteria. For intratumoral infections, frozen stocks were diluted in PBS to infect via intratumoral injections with 5×10^7^ CFU log-phase bacteria. In some instances, intratumoral bacteria was substituted with 100 ug Pam3CSK4 (Invivogen) or supplemented with 50 ug diphtheria toxin (EMD Millipore). Uninfected control animals were mock infected by intratumoral injection of 100 uL PBS. Mice were euthanized 1-20 days after intratumoral injections.

#### Colony forming unit assays from tissues

Tumors were surgically resected following euthanasia, weighed, and transferred to 0.1% NP40 buffer diluted in PBS. Organs were homogenized with a Fisherbrand™ 150 Handheld Homogenizer (Thermo Fisher) and serially diluted on non-TC treated 96-well plates (Genesee). Serial dilutions were plated on BHI + 200 μg/mL streptomycin plates. Plates were incubated overnight at 37°C. The following day, CFUs were counted and normalized by tumor weight.

#### Bone marrow dendritic cell differentiation

Primary BMDCs were generated as previously described using WT B6 mice (day 0).^132^ After differentiation (day 6), BMDC media was replaced with fresh media containing PBS or 1 ug/mL Pam3CSK4 (Invivogen) to stimulate TLR2. 24 hr later (day 7), BMDCs were harvested by transferring media containing nonadherent cells to a centrifuge tube and replacing with an equal volume of cold PBS supplemented with 10 uM EDTA to loosen semi-adherent cells. Nonadherent cells were pelleted by centrifugation, while semi-adherent cells were incubated for 10 min on ice. After incubation, semi-adherent cells were recovered by gentle pipetting, and this suspension was used to resuspend pelleted nonadherent cells. Pooled cells were washed with PBS to remove trace Pam3CSK4. BMDCs were counted and used in Treg co-culture experiments or analyzed by flow cytometry to assess purity and phenotype.

#### Fluorescence-activated cell sorting of Tregs

Treg lineage-trace mice were euthanized and lymph nodes and spleens surgically resected and stored on ice in PBS supplemented with 4 mM EDTA and 2% FBS. Single-cell suspensions of splenocytes and lymphocytes were generated by crushing organs through a 40 um cell strainer (Fisher Scientific) and red blood cells were removed by ACK lysis using RBC lysis buffer (BioLegend). CD4 T cells were magnetically enriched using the EasySep™ Mouse CD4+ T Cell Isolation Kit (STEMCELL Technologies) per the manufacturer’s instructions. Isolated CD4 T cells were stained with fluorescent antibodies in the dark at 4°C for 30 min and washed with PBS prior to resuspension in Opti-MEM (Thermo Fisher) supplemented with 2% FBS for FACS. Naive Tregs (CD4+CD8-CD62L+RFP+GFP+) were sorted with a FACSAria™Fusion cell sorter (BD). Sorting efficiency was determined by post-sort flow cytometric analysis and Treg purity exceeded 95%.

#### Treg-BMDC co-culture assay

FACS-isolated Tregs were mixed with BMDCs or Dynabeads™ Mouse T-Activator CD3/CD28 for T-Cell Expansion and Activation (GIBCO) at a ratio of 4:1::BMDCs:Tregs or 1:3::beads:Tregs. BMDC-Treg cultures were supplemented with 1 ug/mL of Ultra-LEAF™ Purified anti-mouse CD3 Antibody (BioLegend) to mimic TCR signaling. For indirect stimulation of TLR2 on Tregs mediated by DCs, Pam3-stimulated BMDCs were used in some BMDC-Treg cultures. For direct stimulation of TLR2 on Tregs, 1 ug/mL Pam3 (Invivogen) was included in some bead-Treg cultures. All cells were cultured in DMEM supplemented with 10% FBS, sodium pyruvate (GIBCO), L-Glutamine (GIBCO), penicillin-streptomycin (GIBCO), HEPES (GIBCO), MEM Non-Essential Amino Acid Solution (GIBCO), 50 uM B-mercaptoethanol (GIBCO), and 200 IU/mL teceleukin (hIL2; Hoffmann-La Roche). Cultures were incubated at 37°C for 72 hr prior to flow cytometric analysis to assess Treg phenotype and Foxp3 destabilization.

#### Bone marrow chimera generation

Host CD45.1+ mice were subjected to irradiation using an XRad320 (Precision X-Ray Irradiation) on day 0 (550 cGy) and 16 hours later on day 1 (500 cGy). 8 hours after the second dose of radiation, mice received 4×10^6^ bone marrow cells composed of equal parts (2×10^6^ cells/each) *CD45*^.1/.1^;*Tlr2*^+/-^ and *CD45.*^2/.2^;*Tlr2^-^*^/-^ cells via tail vein injection. Bone marrow was collected by surgically resecting femurs and tibias from euthanized mice, cleaning to remove excess tissue, and flushing bone marrow into a 15 mL conical centrifuge tube with a 27-gauge needle using ice-cold RPMI supplemented with 10% FBS, sodium pyruvate (GIBCO), L-Glutamine (GIBCO), HEPES (Gibco), MEM non-essential amino acids (Gibco), penicillin-streptomycin (GIBCO), and 55 uM 2-mercaptoethanol. Red blood cells were removed by ACK lysis using RBC lysis buffer (BioLegend) and remaining bone marrow progenitor cells were washed extensively with ice-cold PBS prior to tail vein injection. Mice received water supplemented with sulfamethoxazole-trimethoprim oral suspension (Ani Pharmaceuticals) for 30 days with daily monitoring following adoptive transfer of bone marrow. Successful reconstitution was assessed by staining tissues with fluorophore-conjugated anti-CD45.1 and -CD45.2 antibodies and flow cytometric analysis. All mice used in experiments were found to have a ∼50:50 ratio of CD45.1+:CD45.2+ cells in all tissues confirming successful reconstitution.

#### Sample preparation for flow cytometry

Flow cytometry was performed on an BD LSR Fortessa X20 (BD Biosciences) or CyTEK Aurora (CyTEK Biosciences) and datasets were analyzed using FlowJo software (Tree Star). Single cell suspensions were prepared as described below (“*Tissue collection and restimulation Assays”*). Dead cells were stained with Live/Dead Fixable Blue Cell Stain kit (Molecular Probes) in PBS at 4°C. Cell surface antigens were stained at 4°C using a mixture of fluorophore-conjugated antibodies. Surface marker stains for murine T cell samples were carried out with anti-mouse CD3 (17A2, BioLegend), anti-mouse CD4 (RM4-5, BioLegend), anti-mouse CD8a (53-6.7, BioLegend), anti-mouse CD25 (PC61, BioLegend), anti-mouse, CD44 (IM7, BioLeged), anti-mouse CD45 (30-F11, BioLegend), anti-mouse CD45.1 (A20, BioLegend), anti-mouse CD45.2 (104, BioLegend), anti-mouse CD62L (MEL-14, BioLegend), anti-mouse CD69 (H1.23F, BioLegend), anti-mouse CD103 (2E7, BioLegend), anti-mouse Lag3 (C9B7W, BioLegend), anti-mouse PD1 (29F.1A12, BioLegend), anti-H-2Kb MuLV p15E Tetramer-KSPWFTTL (MBL), anti-H2-Kb-A2/SIINFEKEL tetramer (NIH tetramer core), and anti-IAb/NEKYAQAYPNVS tetramer (NIH tetramer core) in PBS. Surface marker stains for murine myeloid cell samples were carried out with anti-mouse CD16/32 TruStain FcX (93, BioLegend), anti-mouse CD45 (30-F11, BioLegend), anti-mouse F4/80 (BM8, BioLegend), anti-mouse CD45R (RA3-6B2, BioLegend), anti-mouse CD11c (N418, BioLegend), anti-mouse I-A/I-E (M5/114.15.2, BioLegend), anti-mouse SIRPa (P84, BioLegend), anti-mouse CD11b (M1/70, BioLegend), anti-mouse XCR1 (ZET, BioLegend), anti-mouse CD103 (2E7, BioLegend), anti-mouse Ly6C (HK1.4, BioLegend), anti-mouse Ly6G (1A8, BioLegend), anti-mouse CD14 (Sa14-2, BioLegend), and anti-H2-K(b)-SIINFEKL (25-D1.16, BioLegend) in PBS. Cells were fixed in 4% PFA (Electron Microscopy Sciences) in PBS at room temperature for 15 minutes, permeabilized with 0.1 % Triton X-100 (Thermo Fisher) in PBS at room temperature for 10 minutes, and washed twice with 0.5% BSA (Sigma Aldrich) in PBS prior to intracellular staining. Intracellular staining for murine T cell samples was performed using anti-mouse Foxp3 (FJK-16S, eBioscience) and anti-mouse IL-2 (Dec-16, BioLegend) at 4°C in 0.5% BSA in PBS. Cells were analyzed by flow cytometry the following day to prevent signal loss from fluorescent proteins (mCherry, GFP, tagBFP). Cells were resuspended in PBS and filtered through FlowTubes™ Cap w/ 35 μm Strainer Mesh (Genesee) prior to data acquisition. In some instances, previously published data were reanalyzed to assess the impact of IV+IT *Lm* on the cross-presentation of tumor-derived ovalbumin by APC subsets.^39^

#### Tissue collection and restimulation assays

Tumors and lymphoid organs were surgically resected following euthanasia of mice according to the Institutional Animal Care and Use Committee guidelines of the University of California, Berkeley. Resected tumors were minced to 1 mm^3^ fragments and digested in RPMI media supplemented with 20 mg/mL DNase I (Roche), 125 U/mL collagenase D (Roche), and 2% FBS using an orbital shaker at 37°C for 1 hr. Digested tumors were filtered through a 70 μm cell strainer (Thermo Fisher). Cells from lymphoid organs were prepared by mechanical disruption with the sterile plunger of a 1 mL syringe through a 70 μm nylon mesh cell strainer (Thermo Fisher). Cell suspensions from tumors and lymphoid organs were subjected to ACK lysis to remove red blood cells using RBC lysis buffer (BioLegend) and passed through FlowTubes™ Cap w/ 35 μm Strainer Mesh (Genesee) prior to in vitro stimulation and/or flow cytometric analysis. Simulation was performed with 3-5×10^6^ cells in Opti-MEM media (Thermo Fisher) supplemented with Brefeldin A (eBioscience), 10 ng/mL phorbol 12-myristate 13-acetate (PMA) (Sigma-Aldrich), and 0.25 μM ionomycin (Sigma-Aldrich) at 37°C for 6 hr. Surface staining, fixation, permeabilization, and intracellular staining were performed as described above (“*Tissue collection and preparation for flow cytometry”*).

#### Adoptive transfer experiments

For tumor growth experiments, 2×10^6^ *in vitro* expanded OT-I T cells were transferred IV into *Tlr2^+/-^* and *Tlr2^-/-^* littermate control mice 2 weeks prior to MC38 inoculations. For *in vitro* OT-I expansion, spleens and lymph nodes were collected from OT-I transgenic mice. OT I CD8+ T cells were were activated with 1μg/mL SIINFEKL peptide in DMEM supplemented with 10% FBS, sodium pyruvate (GIBCO), L-Glutamine (GIBCO), penicillin-streptomycin (GIBCO), HEPES (GIBCO), MEM Non-Essential Amino Acid Solution (GIBCO), 55 uM B-mercaptoethanol (GIBCO), and 200 IU/mL teceleukin (hIL2; Hoffmann-La Roche provided by NCI repository, Frederick National Laboratory for Cancer Research). In some experiments where indicated, splenocytes from WT mice immunized 1-2 weeks prior with 10^6^ CFU of *Lm-OVA* IV were expanded identically to OT-Is. In these cases, SIINFEKL/K(b) tetramer staining was performed prior to adoptive transfer to validate that a pure population of T cells (>85% tetramer+) was used for all experiments. For priming experiments, 5×10^5^ naive, CFSE-labeled OT-I T cells were transferred IV into MC38 tumor-bearing WT B6 mice 12 days post tumor inoculation. Naive OT-I T cells were isolated from spleens and lymph nodes using the MagniSort™ Mouse CD8+ T cell Enrichment Kit (Invitrogen) per the manufacturer’s instructions. Isolated naive OT-I T cells were labeled with the CFSE Cell Division Tracker Kit (BioLegend) per the manufacturer’s instructions. CFSE-labeled OT-Is were transferred IV into B6.SJL-*Ptprca Pepcb*/BoyJ (CD45.1) mice 1 day following IT injections of *Lm* or PBS.

#### In vivo antibody-mediated cell depletion

For tumor progression studies, IL-2 depletion was achieved by intraperitoneal injection of 200 μg per mouse of the anti-IL-2 monoclonal antibody clone JES6-1A12 (Leinco, Catalog # I-1042) one day prior IT *Lm* treatment, followed by additional doses twice weekly thereafter.

#### Statistical Methods

p values were obtained from unpaired two-tailed Student’s t tests for statistical comparisons between two groups of values derived from separate mice, or from paired two-tailed Student’s t tests for comparisons between tissues within individual mice, and data were displayed as mean ± SEMs. For multiple comparisons, one-way ANOVA was used. For tumor growth curves, two-way ANOVA was used with Sidak’s multiple comparisons test performed at each time point or by multiple regression analysis p values are denoted in figures by *p < 0.05, **p < 0.01, ***p < 0.001, and ****p < 0.0001.

