## Supplemental figures S1-S8 for "TLR2 signaling uniquely destabilizes tumor Tregs to promote cancer immunotherapy"

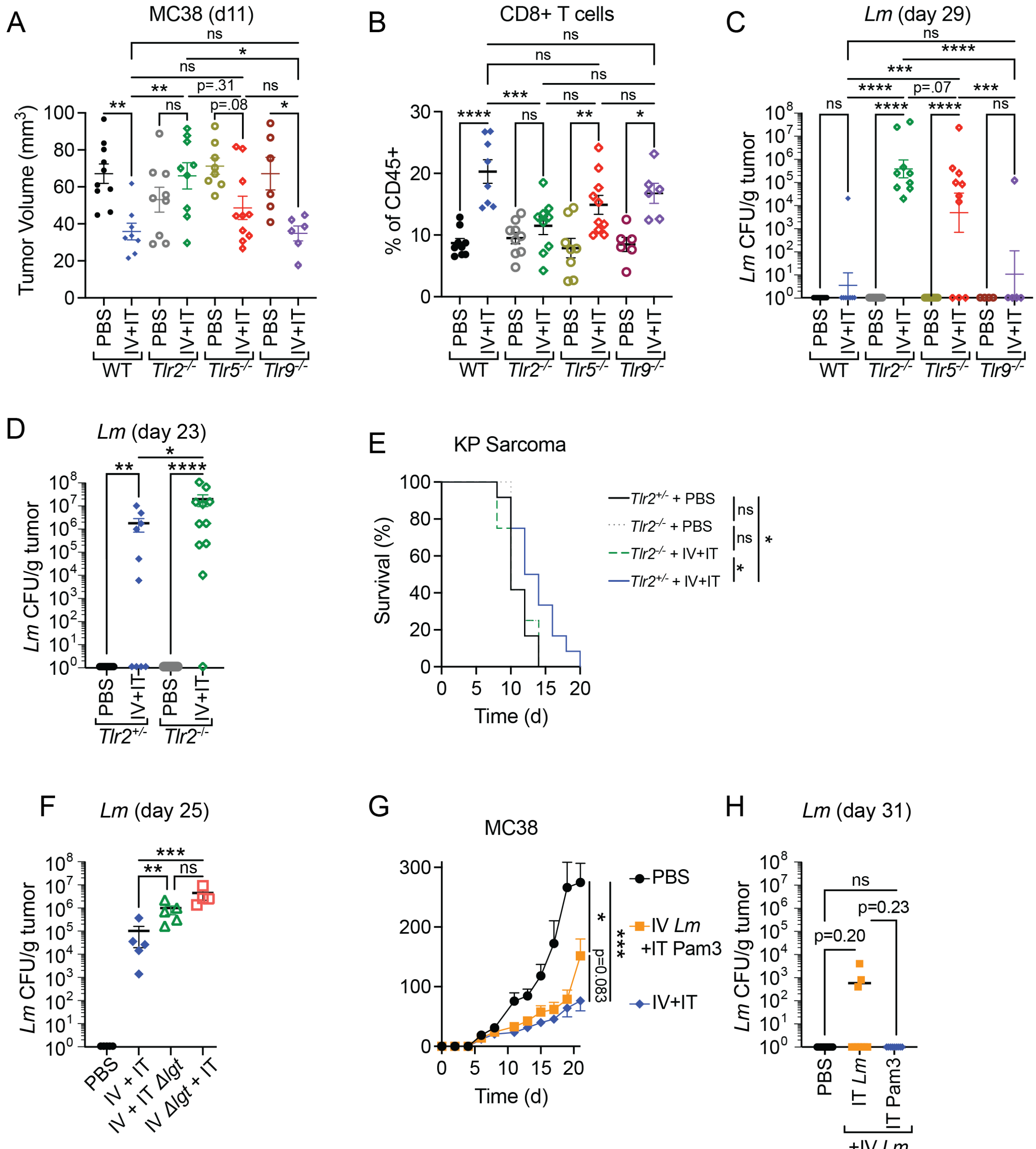


Figure S1. Campbell et al.

**Figure S1 (related to Fig. 1):**

(A) MC38 tumor volume on day 11 of tumor growth from experiment in Figure 1B.

(B) CD8$^{+}$ T cell frequencies in MC38 tumors from experiment in Figure 1B on day 29 of tumor growth.

(C) *Lm* CFUs recovered from MC38 tumors on day 29 of tumor growth in IV+IT *Lm*-treated WT (*Tlr2*$^{+/-}$), *Tlr2*$^{-/-}$, *Tlr5*$^{-/-}$, and *Tlr9*$^{-/-}$ mice from experiment in Figure 1B.

(D) *Lm* CFUs recovered from B16F10 tumors on day 23 of tumor growth in IV+IT *Lm*-treated *Tlr2*+/- or *Tlr2*-/- littermate mice from experiment in Figure 1C.
(E) Percent survival of KP sarcoma-bearing *Tlr2*+/- and *Tlr2*-/- mice receiving IV+IT *Lm* or PBS.
(F) *Lm* CFUs recovered from MC38 tumors on day 25 of tumor growth in IV+IT *Lm*-treated mice that received *LmΔlgt* at the IV stage, IT stage, or *Lm* at both stages of treatment from experiment in Figure 1F.
(G) MC38 tumor growth in IV+IT *Lm*-treated or IV *Lm*+IT Pam3-treated mice up to day 21 from experiment in Figure S1G.
(H) *Lm* CFUs recovered from MC38 tumors on day 31 of tumor growth in IV+IT *Lm*-treated or IV *Lm*+IT Pam3-treated mice from experiment in Figure S1G. Mean ± s.e.m. and one-way ANOVA (A, B, C, D, F, H), log-rank test (E), or two-way ANOVA (G).

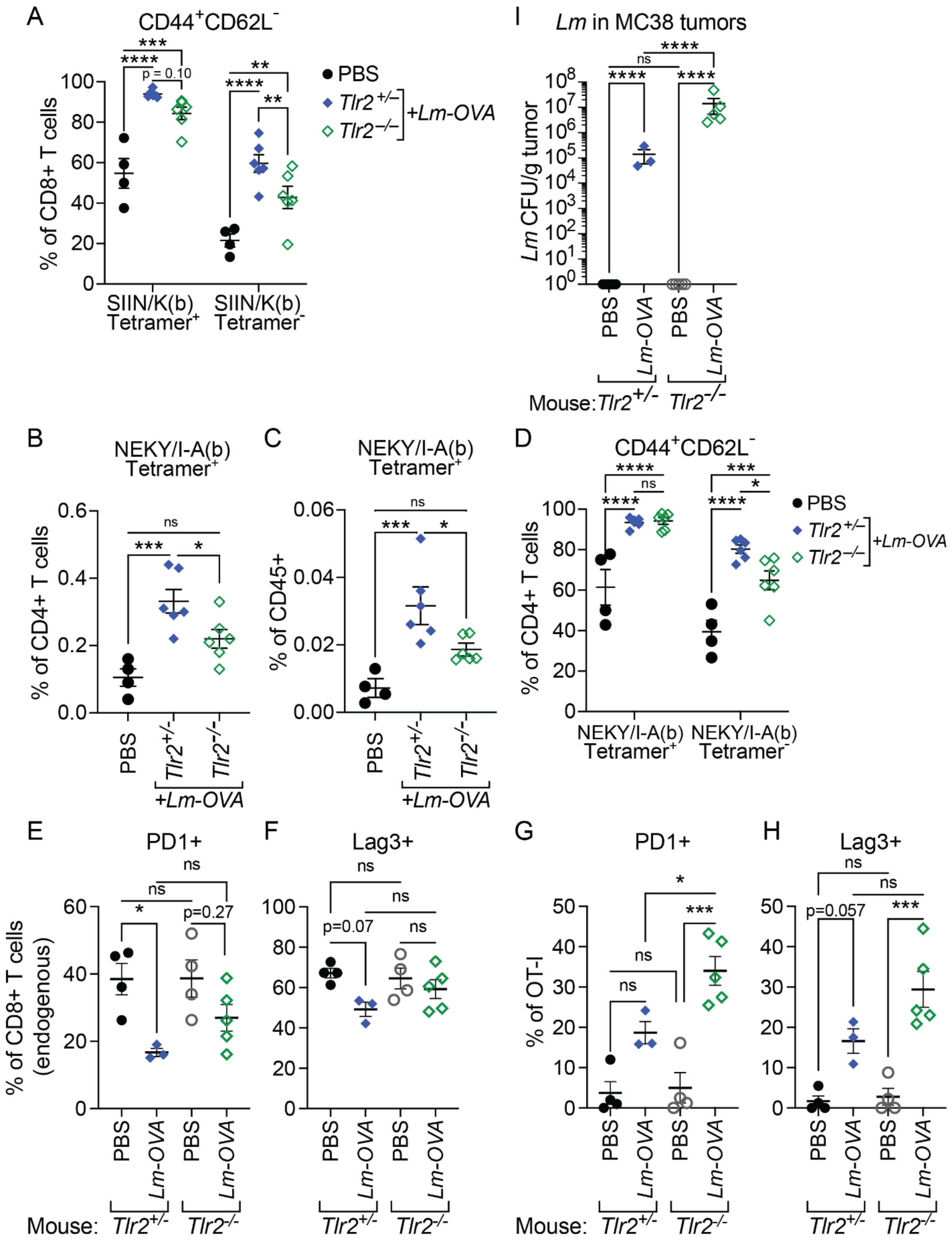


Figure S2. Campbell et al.

**Figure S2 (Related to Fig. 2):**

(A) Frequency of activated CD8+ T cells in the spleen 5 days post-IV *Lm-OVA* in *Tlr2*$^{-/-}$ or *Tlr2*$^{+/-}$ littermate mice stratified by SIIN/K(b)+/-.

(B, C) Frequency of NEKY/I-A(b) tetramer+ CD4+ T cells in the spleens of TLR2$^{-/-}$ and TLR2$^{+/-}$ littermates 5 days post-IV *Lm-OVA* as a percentage of CD4+ T cells (B) or of total CD45+ splenocytes (C).

(D) Frequency of activated CD4+ T cells in the spleen 5 days post-IV *Lm-OVA* in *Tlr2*$^{-/-}$ or *Tlr2*$^{+/-}$ littermate mice stratified by NEKY/I-A(b)+/-.

(E) Expression of PD1 on endogenous CD8 T cells form tumors of *Tlr2*$^{-/-}$ or *Tlr2*$^{+/-}$ littermate mice treated with PBS or IV+IT *Lm* on day 31 (20 days post-IT *Lm*).

(F) Expression of Lag3 on endogenous CD8 T cells form tumors of *Tlr2*$^{-/-}$ or *Tlr2*$^{+/-}$ littermate mice treated with PBS or IV+IT *Lm* on day 31 (20 days post-IT *Lm*).

(G) Expression of PD1 on OT-I T cells form tumors of *Tlr2*$^{-/-}$ or *Tlr2*$^{+/-}$ littermate mice treated with PBS or IV+IT *Lm* on day 31 (20 days post-IT *Lm*).

(H) Expression of Lag3 on OT-I T cells form tumors of *Tlr2*$^{-/-}$ or *Tlr2*$^{+/-}$ littermate mice treated with PBS or IV+IT *Lm* on day 31 (20 days post-IT *Lm*).

(I) *Lm-OVA* CFUs recovered on day 31 from the tumors of *Tlr2*$^{-/-}$ or *Tlr2*$^{+/-}$ littermate mice.. Mean ± s.e.m. and one-way ANOVA (B, C, E, F, G, H, I) or two-way ANOVA (A, D).

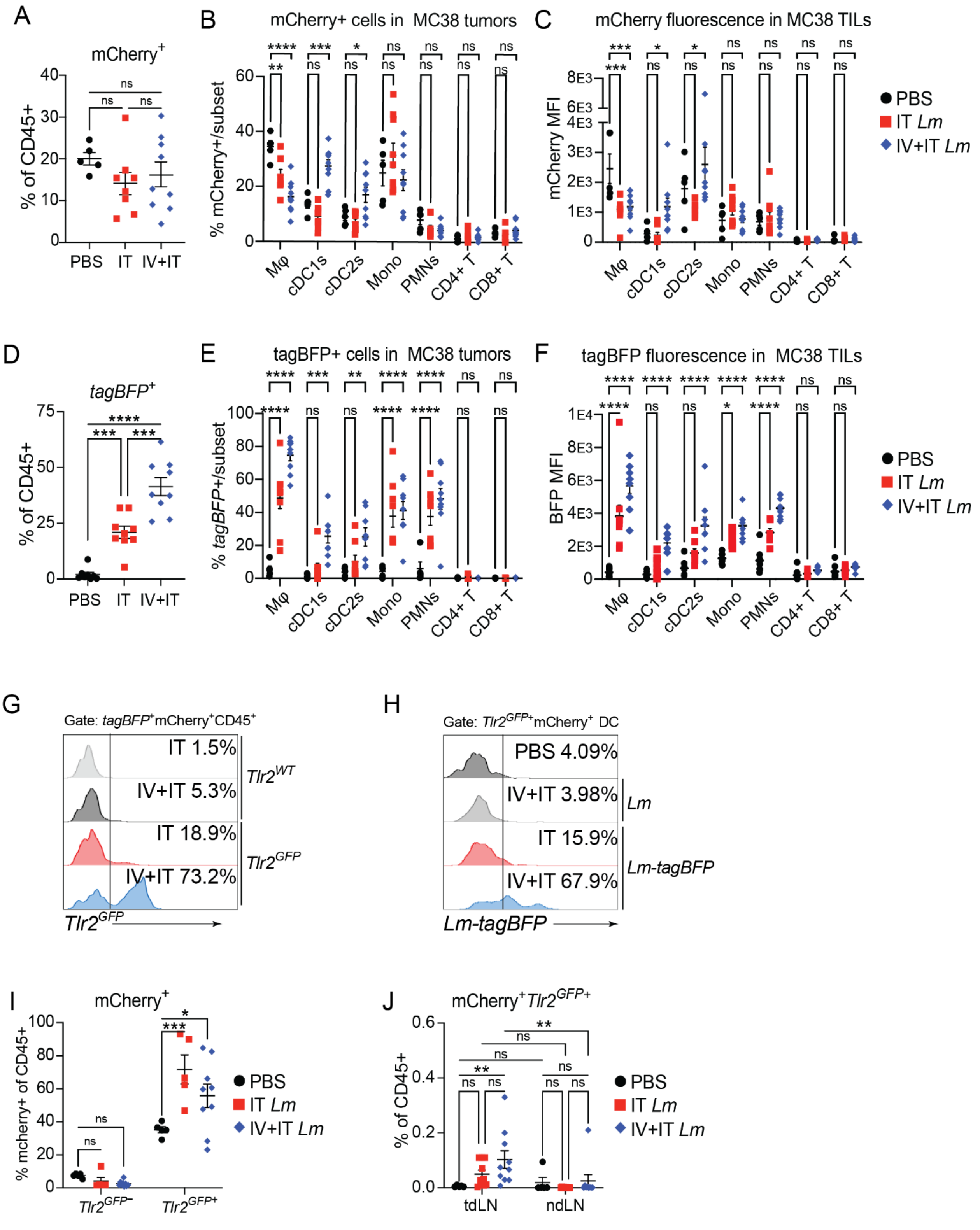


Figure S3. Campbell et al.

**Figure S3 (related to Fig. 3):** All data from MC38 tumors 24 hr post-IT *Lm-tagBFP* or PBS (day 12).
(A) Total mCherry+CD45+ cell frequency.
(B) mCherry+ cell subset frequency.
(C) MFI of mCherry in CD45+ cell subsets.
(D) Total *tagBFP*+CD45+ cell frequency.
(E) *tagBFP*+ cell subset frequency.
(F) MFI of *Lm-tagBFP* in CD45+ cell subsets.
(G) Representative flow cytometry histograms showing *Tlr2*$^{GFP}$ fluorescence within *tagBFP*+mCherry+ CD45+ cells in MC38 tumors of *Tlr2*$^{GFP}$ or WT control mice treated with IT only or IV+IT *Lm-tagBFP*.
(H) Representative flow cytometry histograms showing *tagBFP* fluorescence within *Tlr2*$^{GFP}$+mCherry+ DCs in MC38 tumors treated with PBS, IV+IT *Lm*, IT *Lm-tagBFP*, or IV+IT *Lm-tagBFP*.
(I) mCherry+ cell frequency among *Tlr2*$^{GFP}$- and *Tlr2*$^{GFP}$+ cells in MC38 tumors.
(J) mCherry+*Tlr2*$^{GFP}$+ cell frequency in lymph nodes 24 hr post-IT *Lm*. Mean ± s.e.m. and one-way ANOVA (A, D) or two-way ANOVA (B, C, E, F, I, J).

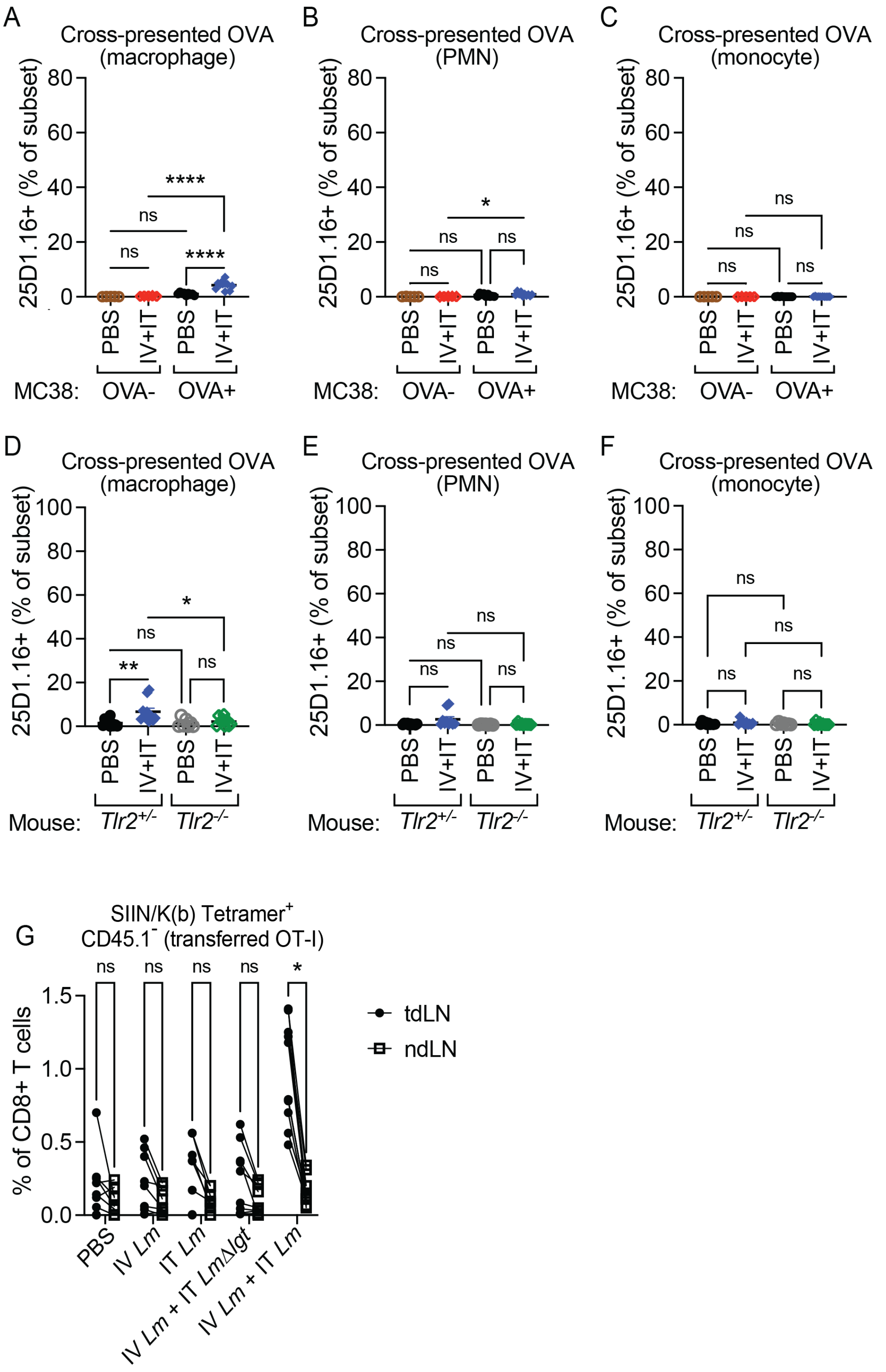


Figure S4. Campbell et al.

**Figure S4 (related to Fig. 4)**:
(A, B, C) Frequency of SIIN/K(b)+ cells among macrophages (A), PMNs (B), and monocytes (C) detected by 25D1.16 staining in tumors of MC38-B2m$^{KO}$ control or MC38-B2m$^{KO}$-OVA$^{+}$ tumor-bearing mice.
(D, E, F) Frequency of SIIN/K(b)+ cells among macrophages (D), PMNs (E), and monocytes (F) detected by 25D1.16 staining in tumors of MC38-B2m$^{KO}$-OVA$^{+}$ tumor-bearing *Tlr2*$^{+/-}$ and *Tlr2*$^{-/-}$ littermate mice.
(G) Paired analysis of the frequency of transferred OT-I among total CD8 T cells in the tdLN and ndLN. Mean +/- s.e.m. and one-way ANOVA (A, B, C, D), paired Student's t-test (E).

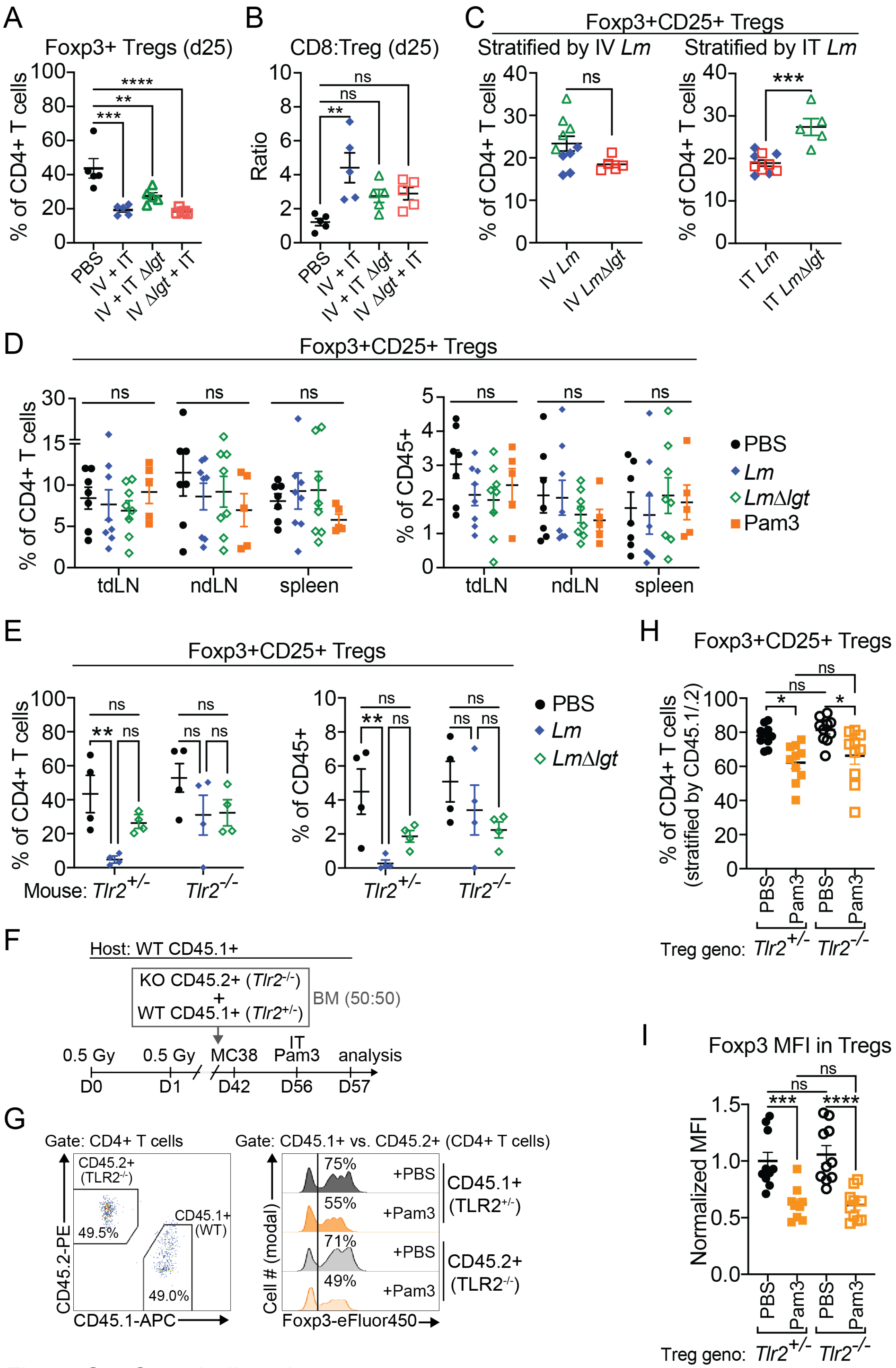


Figure S5. Campbell et al.

**Figure S5 (related to Figures 5 and 6):**

(A) Frequency of Tregs among CD45+ cells on day 25 of tumor growth in mice treated with IV+IT *Lm* using *LmΔlgt* at the IV or IT stage of treatment.

(B) Ratio of $CD8^+$ T cells:Tregs on day 25 of tumor growth in mice treated with PBS, IV+IT *Lm*, or IV+IT *Lm* using *LmΔlgt* at the IV or IT stage of treatment.

(C) Frequency of Tregs among $CD4^+$ T cells cells on day 25 of tumor growth in mice treated with IV+IT *Lm* using *LmΔlgt* at the IV or IT stage of treatment stratified by IV (left) or IT (right).

(D) Frequency of Tregs as a % of $CD4^+$ T cells (left) or of total $CD45^+$ lymphocytes/splenocytes (right) in the tdLN, ndLN, and spleen 24 hr post-IT treatment.

(E) Frequency of Tregs as a % of $CD4^+$ T cells (left) or of total $CD45^+$ TILs (right) in the tumors of *Tlr2*$^{+/-}$ or *Tlr2*$^{-/-}$ littermates 24 hr post-IT *Lm*.

(F) Experimental outline for generating *Tlr2*$^{+/-}$;$CD45^{.1/.1}$/*Tlr2*$^{-/-}$;$CD45^{.2/.2}$ mixed bone marrow chimera mice and testing whether Treg-intrinsic TLR2 signaling mediates the reduction in tumor-infiltrating Treg frequency following IT Pam3.

(G) Representative flow cytometry histograms showing the gating of CD45.1+ (*Tlr2*$^{+/-}$) and $CD45.2^+$ (*Tlr2*$^{-/-}$) $CD4^+$ T cells in MC38 tumors and the decrease in both the frequency $Foxp3^+$ cells and intensity of Foxp3 expression in both compartments 24 hr post-IT Pam3 or PBS.

(H). Quantification of $Foxp3^+CD25^+$ Tregs as a % $CD45.1^+$ or $CD45.2^+$ $CD4^+$ T cells in MC38 tumors.

(I) Quantification of Foxp3 MFI within $CD45.1^+$ or $CD45.2^+$ Tregs. Mean +/- s.e.m. and one-way ANOVA (A, C, D, G, H) or Student's t-test (B).

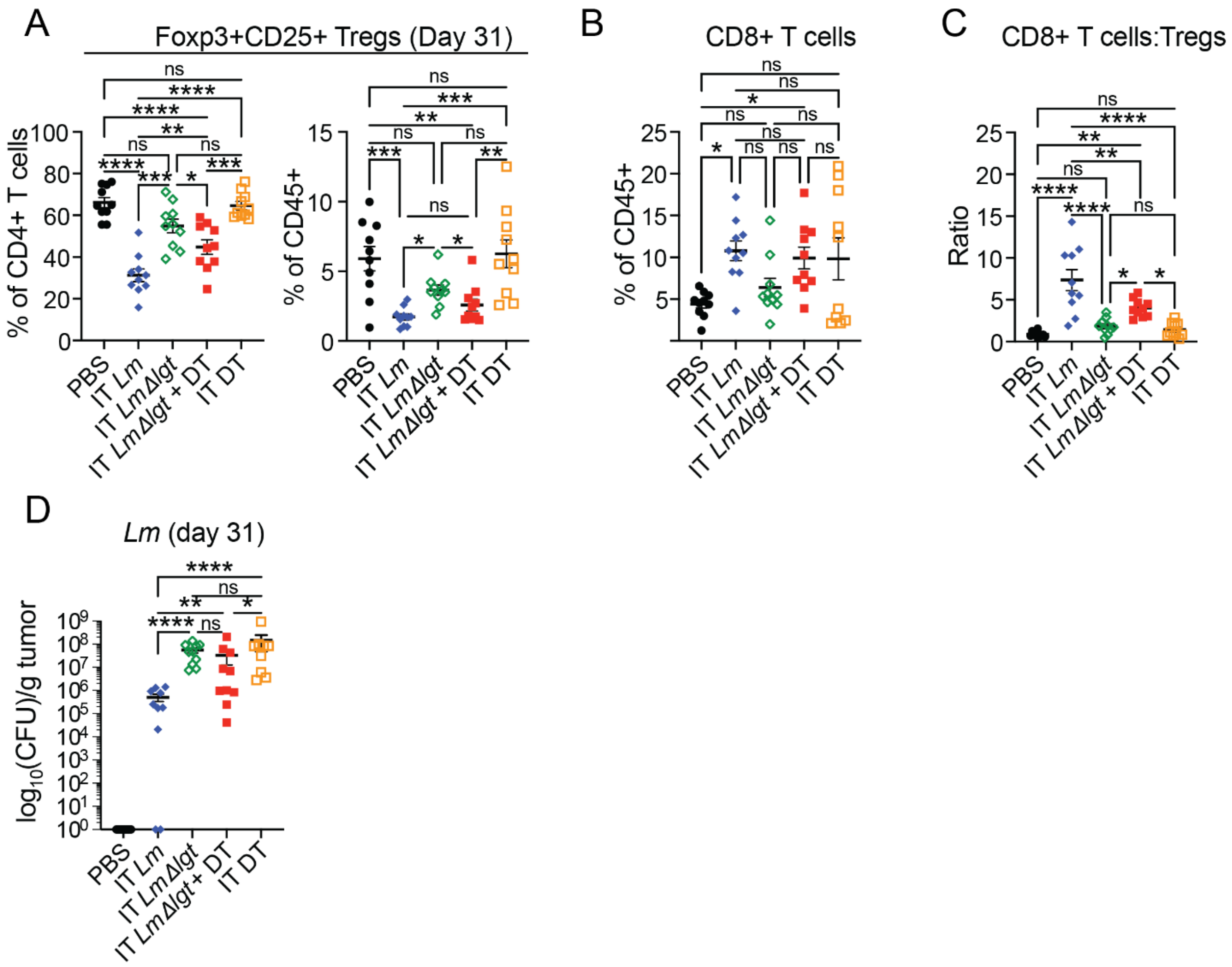


Campbell *et al.*, Figure S6

**Figure S6 (related to Fig. 7A-B)**:
(A) Frequency of Tregs as % of CD4+ T cells (left) or of total CD45+ TILs (right) on day 31 of tumor growth following IV *Lm* + IT treatment with indicated modifications.
(B) Frequency of CD8+ T cells on day 31 of tumor growth.
(C) Ratio of CD8+ T cells:Tregs on day 31 of tumor growth.
(D) *Lm* CFUs recovered from tumors treated with *Lm* IV+IT in the indicated conditions on day 31 of tumor growth. Mean +/- s.e.m. and one-way ANOVA (A, B, C, D).

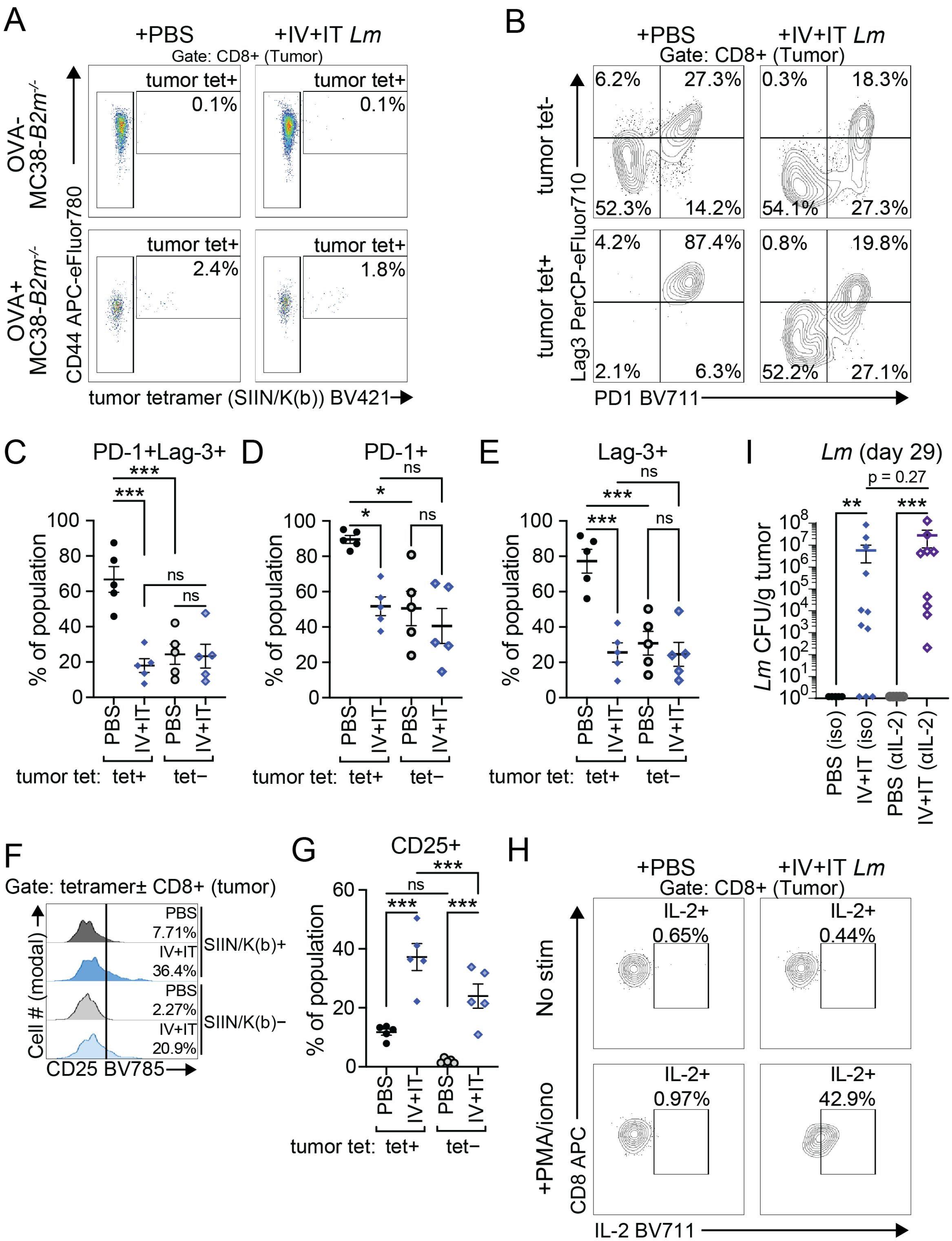


Campbell *et al.*, Figure S7

**Figure S7 (related to Fig. 7C-G)**:
(A) Representative flow cytometry plots showing the expansion of tumor-specific SIIN/K(b) tetramer+ CD8+ T cells in MC38-B2m$^{KO}$-OVA± tumors 1 day post-IT *Lm* or PBS (see Figure 4A).
(B) Representative flow cytometry plots showing Lag3 and PD1 staining in tumor-specific SIIN/K(b) tetramer± CD8 T cells in PBS or IV+IT *Lm*-treated OVA$^{+}$ tumors 24 hr post-IT.
(C) Co-expression of PD-1 and Lag3 on tumor-specific and bulk CD8+ T cells.
(D) Expression of PD-1 on tumor-specific and bulk CD8+ T cells.
(E) Expression of Lag3 on tumor-specific and bulk CD8+ T cells.
(F) Representative flow cytometry histograms showing CD25 staining in SIIN/K(b) tetramer± CD8 T cells in PBS or IV+IT *Lm*-treated OVA$^{+}$ tumors 24 hr post-IT.
(G) Expression of CD25 on tumor-specific and bulk CD8+ T cells
(H) Representative flow cytometry plots showing IL-2 staining in CD8 T cells from PBS or IV+IT *Lm*-treated tumors 7 days post-IT after 6 hr restimulation with media or PMA+ionomycin.
(I) *Lm* CFUs recovered from IL-2-depleted or control tumors on day 29 of tumor growth. Mean +/- s.e.m. and one-way ANOVA (C, D, E, G, I).

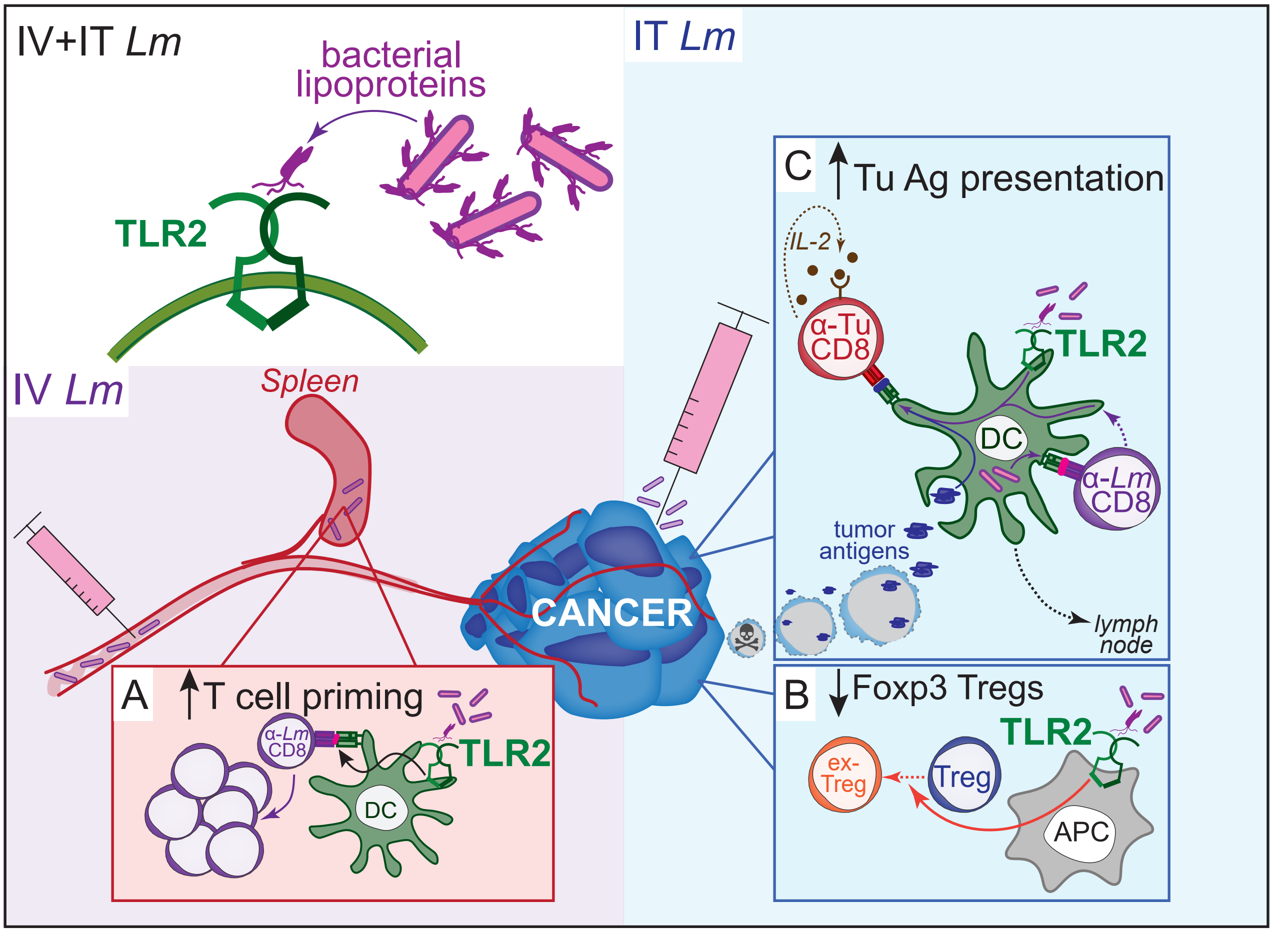


**Figure S8 (graphical abstract): *Lm* lipoproteins mediate TLR2-dependent responses to IV+IT *Lm*.**

(A) TLR2 contributes to optimal priming of anti-*Lm* T cells.

(B) APCs stimulated by TLR2 drive Foxp3 downregulation in tumor-infiltrating Tregs to generate ex-Tregs.

(C) Reduced Treg immunosuppression and direct TLR2 stimulation of DCs drive tumor antigen uptake and cross-presentation to promote tumor-specific CD8 T cell function and and IL-2 signaling.
